# Real-time contextual feedback for closed-loop control of navigation

**DOI:** 10.1101/473108

**Authors:** Judith Lim, Tansu Celikel

## Abstract

**Objective:** Close-loop control of brain and behavior will benefit from real-time detection of behavioral events to enable low-latency communication with peripheral devices. In animal experiments, this is typically achieved by using sparsely distributed (embedded) sensors that detect animal presence in select regions of interest. High-speed cameras provide high-density sampling across large arenas, capturing the richness of animal behavior, however, the image processing bottleneck prohibits real-time feedback in the context of rapidly evolving behaviors.

**Approach:** Here we developed an open-source software, named PolyTouch, to track animal behavior in large arenas and provide rapid close-loop feedback in ~5.7 ms, ie. average latency from the detection of an event to analog stimulus delivery, e.g. auditory tone, TTL pulse, when tracking a single body. This stand-alone software is written in JAVA. The included wrapper for MATLAB provides experimental flexibility for data acquisition, analysis and visualization.

**Main results:** As a proof-of-principle application we deployed the PolyTouch for place awareness training. A user-defined portion of the arena was used as a virtual target; visit (or approach) to the target triggered auditory feedback. We show that mice develop awareness to virtual spaces, tend to stay shorter and move faster when they reside in the virtual target zone if their visits are coupled to relatively high stimulus intensity (≥49dB). Thus, close-loop presentation of perceived aversive feedback is sufficient to condition mice to avoid virtual targets within the span of a single session (~20min).

**Significance:** Neuromodulation techniques now allow control of neural activity in a cell-type specific manner in spiking resolution. Using animal behavior to drive closed-loop control of neural activity would help to address the neural basis of behavioral state and environmental context-dependent information processing in the brain.

## 1. Introduction

Animal navigation is a product of closed-loop neural computations [1]. The brain computes a motor plan based on not only the sensory information it gathers from the world but also its expectations given the animal’s previous experience and task-relevant requirements. For instance, a mouse navigating through a maze will alter its trajectory given its experience during prior exposures to the same arena [2–5], as motor strategies driving navigation are continuously shaped by external and internal signals [6–8]. Traditionally this closed-loop control process is studied as an “open-loop” using a stimulus-response design, where a stimulus with a fixed temporal pattern evokes a behavioral or neural response that is typically analyzed offline. This provides only a correlative understanding of the inherent close-loop neural computations that underlie behavior. Instead, to causally address the neural circuits that give rise to dynamic behaviors, e.g. navigation, one needs to interfere with neural activity in the context of ongoing behavior.

Recent technological advancements have enabled researchers to close the loop [9,10] with real-time experimental control of neural [11–14] and behavioral signals [15–19]. In particular, closed-loop optogenetic neural control has provided insight into cell-type-specific neuronal population dynamics in relation with seizure control [14,20,21], sensory perception [22], and spatial navigation [23]. Nevertheless, the real-time monitoring of behavior remains a challenging task for many researchers, because it demands (1) reliable event-detection at a high temporal resolution in real-time, (2) low-latency communication with peripheral devices to trigger feedback and (3) flexibility in terms of software and hardware integration. Previous animal studies have commonly used a single or array of infrared (IR) beam sensors to detect simple motion events (e.g. entering an area, lick in a reward port, [24] or a grid of sensors, including pressure sensors [25], to monitor locomotion with a delay in the millisecond range [2,3,5,26–28]) to overcome most of these limitations. Although these systems are fast and reliable, their spatial resolution is heavily limited by the number of sensors deployed. Despite the availability of various other sensors, including microwave based motion detectors [29], ultrasonic microphones [30,31], radiofrequency detectors [32], global positioning systems [33], and heat (infrared) sensors [34], majority of existing tracking systems rely on video cameras, as they provide detailed images of whole bodies [4,35–38],[39], individual limbs [40–42], face and whisker motion [43], and eye movements [44,45]. A major drawback, however, is that behavioral classification requires several image processing steps to detect and identify the object of interest which takes several tens of milliseconds using the current state-of-art algorithms and standard computing infrastructure. This typically prevents rapid, i.e. in the range of milliseconds, closed-loop feedback. As an alternative, other studies have employed a hybrid touch-based imaging approach to counter the speed bottleneck of image processing pipelines [46–48]. In this approach, the animal motion is tracked on a transparent surface, where contact points cause scattering of IR light that is captured with a camera. This produces high-contrast images and requires few image processing steps, and thus enables faster image processing.

The utility of touch-based methods in prior studies has not yet been generally extended to multi-touch interfaces [49],[50], in particular for rapid close-loop feedback applications. Since the introduction of accessible multi-touch sensing technology in 2005 [51], most applications targeted human-computer interactions, including using mobile phones, tablets and virtual reality systems [52]. For this very reason, all modern computers and mobile computing devices come with built-in controller drivers for touch sensors that automatically recognize touch input. We extended the utility of this multi-touch technology to develop PolyTouch, an open-source software to track animal behavior in an open field using an IR sensor frame and triggered close-loop feedback with a delay of ~5.7 ms, i.e. between behavioral event detection and stimulus delivery, while continuously digitizing animal (paws and/or body) location at 156.2 Hz. The tracking software is multi-touch capable. It includes a GUI which provides users online measures of the spatial position, traveled distance, body speed, heading direction, relative distance to any user-defined (virtual) target and basic behavioral states. The user can flexibly create any close-loop stimulus protocol that depends on locomotion variables accessible in the output file. We further provide a wrapper in MATLAB for easy integration of PolyTouch in data acquisition and analysis pipelines.

As a proof-of-principle, we tested our closed-loop system in two different place awareness paradigms. First, to provide discrete (positional) feedback, PolyTouch was used to deliver tone pulses whenever a mouse entered a user-defined virtual target zone in an open field arena. We found that the animal tended to stay shorter and moved faster in the target zone if the sound intensity was relatively high (>49 dB), implicating that the animal was aware of the virtual zone. In our second paradigm, a continuous (distance) feedback tone was triggered with a frequency that scaled with the animal’s position relative to a virtual target zone with either increasing or decreasing frequencies (range: 150 - 750 Hz). We observed that the animal adapted its exploration to maximize time spent in the portion of the arena where the frequency was lowest. This approach for animal tracking and rapid sensory feedback with a latency of ~5.7 ms between behavioral event detection and stimulus delivery could prove useful for a broad range of systems neuroscientists studying the principles of behavior, including but not limited to the generation of sensation, perception, action, circuits of learning and memory among others.

## 2. Materials and Methods

### 2.1. Animals

Adult transgenic mice (N = 3), B6;129P2-Pvalbtm1(cr)Arbr/J were bred locally and maintained under *ad libitum* access to food and water. Animals were socially housed on a 12h light/dark cycle. Experimental procedures have been performed in accordance with the European Directive 2010/63/EU, guidelines of the Federation of European Laboratory Animal Science Associations, and the NIH Guide for the Care and Use of Laboratory Animals, and approved by an institutional ethics committee.

### 2.2. Animal tracking and close-loop instrumental control with PolyTouch

PolyTouch is an open-source software written in JAVA that enables animal tracking with real-time elementary behavioral classification while providing rapid feedback at millisecond resolution (Figure 1A-B). The software consists of tracking and feedback modules, the source code is made available via GitHub (https://github.com/DepartmentofNeurophysiology/PolyTouch). The user can run the software as a standalone program or call it in MATLAB (Mathworks) environment. It can be deployed in conjunction with any touch input device, let it be a touch screen with a fixed screen resolution, mouse, touch pen or any interface that utilizes USB Touchscreen Controller (Universal) driver. In this study, we used a standard notebook computer (iCore 5 Intel processor, 8GB DDR2 RAM) and a custom infrared touch frame (wavelength = 850 nm) with 4096 pairs of infrared emitters and detectors placed orthogonally to each other along the four edges of a frame (91 × 52.6 cm; spatial resolution = 0.050 cm). The frame was placed around an arena made of plexiglass (90.5 × 52.6 cm; plexiglass thickness = 1 cm) placed in a sound isolation chamber (Figure 1A). The setup was illuminated by two strips of white light emitting diode arrays placed on the chamber ceiling. Two sound boxes (Speedlink USB speakers, Jöttenbeck GmbH, Germany) were placed on the right- and left half of the arena to provide auditory positional feedback in two proof-of-principle place awareness paradigms (Figure 1A; Section 2.4). The spatial resolution of tracking is scaled by the monitor size of the DAQ computer monitor (23.6 inches, 1920 × 1080 pixels), but is determined by the spatial resolution of the IR sensor frame (i.e. 0.050 cm). Commercially, IR sensor frames are sold as “IR touch over-frame”, “touch frames”, “multi-touch frames” by various manufacturers.

The tracking algorithm makes use of open source jni4net (https://github.com/jni4net/jni4net/) and JWinPointer libraries (http://www.michaelmcguffin.com/code/JWinPointer/) to read the X and Y coordinates of simultaneously detected contact points (Figure 1B). Alternative libraries, e.g. Simple Multi-Touch Toolkit (SMT) (https://github.com/vialab/SMT), could be used instead of JWinPointer to provide cross-platform support. Depending on the relative distance between the ground and the sensor, different portions of the body including limbs, tail, and head can be detected. For the experiments described herein (Section 2.3-4), the sensor was placed on a flat plexiglass sheet aimed for limb detection. It performs behavioral classification based on the temporal and directional changes in body motion. The user interface displays the current animal position (in X,Y), center-of-mass (COM) position, elapsed time (in s), distance traveled (in cm), body speed (in cm/s), the relative position of the (virtual) target (in cm), and basic behavioral states during data acquisition (Figure 1A). Behavioral state identification was based on motion profiling of the animal and included discrete states of animal “moving” (body speed > 1 cm/s), “immobile” (body < 1 cm/s), and advancing “on the ground” or “off the ground”, as the body part moves out of the 2D detection/sensor plane. Simple behavioral state classification allows the user to monitor animal behavior, which could also be used to trigger stimulus feedback.

**Figure 1.**
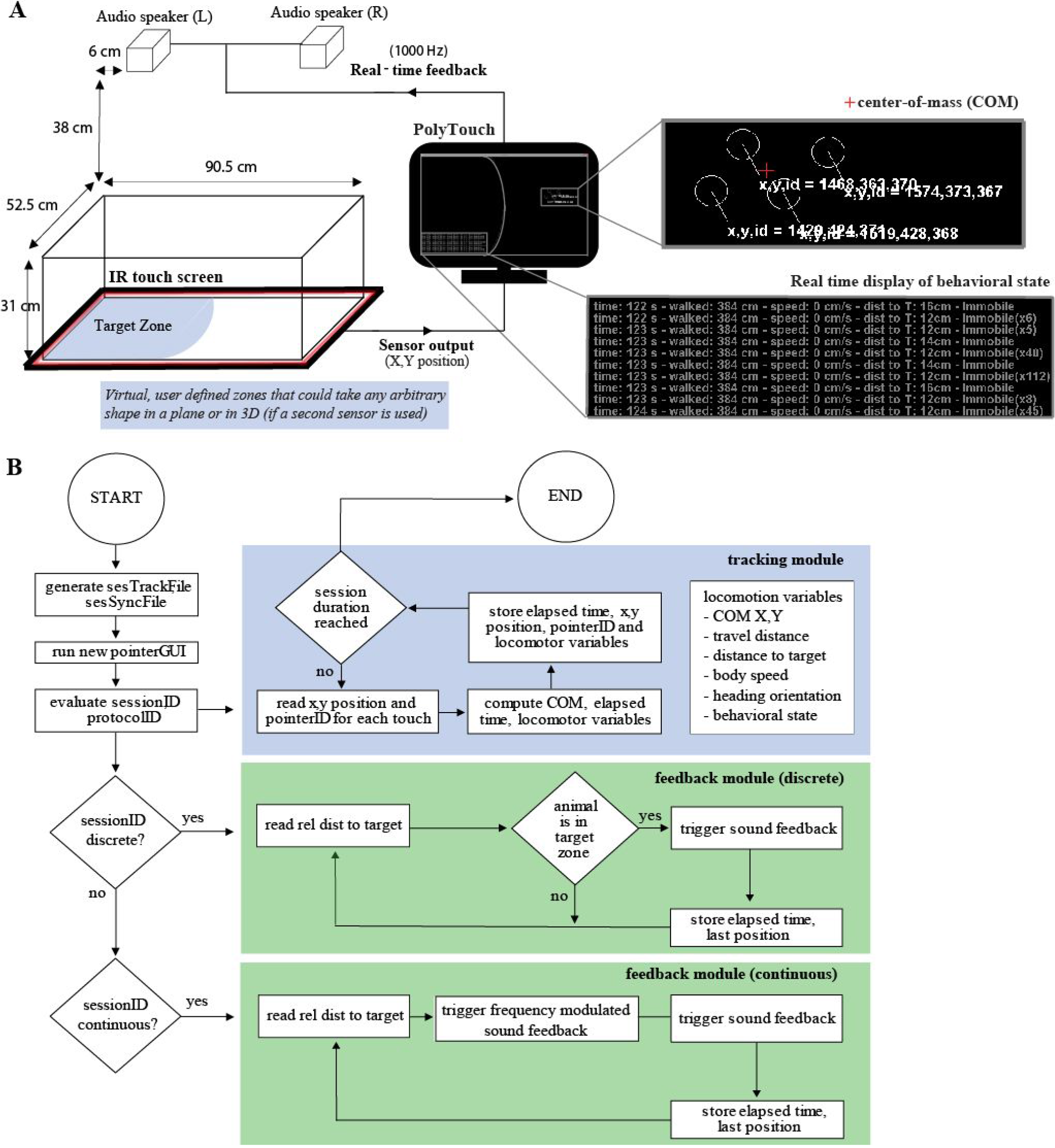
Experimental setup. **(A)** Schematic representation of the experimental setup. PolyTouch tracks animal position and provides control signals for real-time stimulus feedback. The stimulus delivery can be gated by animal behavior and animal’s approach or entry to select regions of interest called virtual target zones. Snapshots of the real-time feedback to the user are shown on a black background. A randomly selected epoch where an animal travels along a path in an experiment is shown. Detected single touches (white circles) along with the calculated center of mass (COM, red cross) provide location information. Graphical User Interface (GUI) also provides real-time information on behavioral state feedback elapsed time (s), travel distance (cm), body speed (cm/s) and distance to the target zone (cm). **(B)** State-flow diagram of the PolyTouch.

PolyTouch assigns a unique ID to each simultaneously detected touch point (e.g. id 1, 2, 3 for paw 1, 2, 3) and updates the touch information (X,Y position, event state, etc.) that is coupled to that particular ID until the touch point is out of range of the sensor, currently at a (sampling) rate of ~156.2 Hz. This means that different events over time can be registered and coupled to the same touch (e.g. paw 1 stops moving: event state changes from “mobile” to “immobile”). In turn, same events can be registered for different touches and thus different touch IDs (e.g. paw 1 and 2 stop moving: event state changes to “immobile” for the two touches). The touch events are updated and exported to an ASCII file as comma separated values at an average rate of 156.2 Hz (± s.d. = 3.5 Hz). Because the data export is restricted to only those time points when the animal is in motion, the file size is kept to a minimum (1 min of behavioral data is ~ 79 KB). The sampling rate of the animal position depends on the CPU load of the data acquisition computer as well as the serial processing of each simultaneous touch events, a constraint of the JAVA architecture. The spatial location in a 2D plane (in pixels) and timestamp (in nanoseconds) of each event are stored along with the device identity (device ID, useful in case multiple sensors are in tandem), touch event identity (touch ID), behavioral state (mobile, immobile, on the ground, off the ground), travelled distance (in cm) and body speed (in cm/s). Timestamps were computed based on the current value of the system timer at nanosecond resolution relative from the moment tracking motion was initiated by the user. To ensure that the motion of the computer mouse does not interfere with the tracking, the computer mouse is assigned to the touch ID of 1 and ignored during the rest of the processing. Note that PolyTouch is not designed for notebook computers (laptops) with touch screen displays whose resolution can be adjusted with multi-touch gestures.

The feedback module controls stimulus delivery in respect to the animal position and context as determined by the user, and generates an external ASCII file to store the timestamps of triggered feedback events (in nanoseconds) that could be used to measure the cycle speed (Figure 1B). The feedback latency is defined as the time delay between the detection of a behavioral event and stimulus delivery. The tracking and feedback modules of PolyTouch run in parallel. Therefore temporal delays associated with digitization of the animal location by the tracking module, i.e. ~6.4 ms corresponding to sampling at ~156.2Hz, and the feedback latency, i.e. on average ~5.7 ms (see Supplemental Figure 1) are independent of one another and do not linearly sum. Considering the slowly evolving nature of animal navigation, the primary temporal constraint for the speed of feedback is the temporal delay introduced by the processing performed by the “feedback module”.

We used the feedback module to deliver auditory stimuli by generating an analog output using the speaker port of the computer, however, it could also be used to communicate with other hardware using analog signals, including TTL pulses. The feedback module continuously reads the X,Y position of the last stored touch event from the external file (write latency: ~1.01 ms, read latency: ~0.4 ms for an HDD spinning at 5400 rpm) and its speed depends on several processing times including behavioral classification (i.e. ~0.001 ms), motion parameters calculation (e.g. COM, body speed; i.e ~0.075 ms), stimulus generation and delivery (i.e. 4.6 ms); all values are measured using a standard mid-range notebook computer (iCore 5 Intel processor, 8GB DDR2 RAM). The feedback is delivered based on either user-defined scenarios or calculated measures of ongoing animal movement, providing real-time feedback to the animal, e.g. in respect to its relative distance to (virtual) target. In this study we deployed two scenarios: **discrete (positional) feedback** that delivers auditory stimulus with a principal frequency of 150 Hz (for the frequency spectrum; see Supplemental Figure 2) whenever the animal is in the user-defined virtual target zone; **continuous (distance) feedback,** which delivers a frequency modulated tone that scales with the relative distance of the animal to the virtual target zone. PolyTouch is terminated when the user-specified session duration is reached or can be interrupted anytime if the user closes the graphical user interface (GUI).

### 2.3. Evaluation of the tracking performance of Polytouch

To estimate the error in tracking performance of PolyTouch, we performed two different controls. First, we tracked a computer controlled robotic ball (Sphero 2.0 ORBOTIX 3.81 cm radius) using two identical, independent sensor frames (61.5 × 35.8 cm). The frames were stacked on top of each other (with 2 cm in between) and each connected to a separate computer to simultaneously sample the position of the robot independently. To minimize the IR light scattering from the beam emitters of the other device, we rotated the top frame 180 degrees clockwise relative from the bottom frame. The robot was programmed to move according to a predefined set of heading directions (0 to 360 deg; total n directions = 72566) with a random speed between 0 to 20 cm/s for a period of 15 minutes. The given heading directions were obtained from a behavioral session recorded previously (i.e. baseline session of mouse #1). As a separate control, we compared the tracking performance of PolyTouch against videographic tracking of mouse location. Manual offline tracking was performed in videos recorded at 30 fps in 2MP resolution (Sony IMX322 sensor, ELP FHD06H) using a custom-written script in MATLAB. An experienced human operator tracked a region selected in the back of the animal in every frame.

### 2.4. Place awareness task with virtual targets

To establish place awareness of virtual target zones, animals were placed in an open field where spatial auditory feedback was provided in two distinct paradigms. In paradigm one, animals (N = 2) received discrete auditory feedback based on the animal’s position in respect to the target zone (number of sessions = 5). In the first session, animals could freely explore the arena for 6 minutes in the absence of any auditory stimulation (0 dB; baseline condition). During the following three sessions, the feedback was provided whenever the animal entered the target zone. The sound intensity was increased across sessions (39 - 49 - 59 dB; low - intermediate - high tone condition, respectively; 15-20 min per session). The target zone location within the arena and the order of the sound intensity were randomly assigned between sessions and animals. In the final session, the tone (3x 39, 49, 59 dB; 10 s each; session duration = 15-20 min) was presented pseudorandomly, independent from the animal location, to control for the sound induced changes in animal mobility. In paradigm two, an animal (N = 1) was first placed in the open field with no auditory feedback (0 dB; baseline condition). In two subsequent sessions, continuous feedback was delivered as a frequency-modulated tone that scaled with the animal’s distance to a virtual target zone with increasing and decreasing frequencies respectively (frequency steps: 150-300-450-600-750 Hz). The open field was cleaned with ethanol (70%) after each session. The inter-session interval was 7 min until after the 3rd session. Afterward, the interval increased to 5 and 10 days for the remaining two sessions in the protocol.

### 2.5. Data analysis

#### 2.5.1. Behavioral analyses

To assess locomotion, the body position was computed as the center-of-mass (COM) of multi-touch events for each time point. The body position was resampled at 200 Hz by averaging samples within non-overlapping but consequential 5 ms bins in 2D (X,Y). If no samples fell within a bin (i.e. no motion was detected), the previous X,Y value was assigned. The resulting matrix was used to quantify the mobility duration (s), body speed (cm/s), and body direction (deg) as a function of time (s) and body position (X,Y).

Mobility duration was represented as a spatial density map and computed as the time spent in a given location by dividing the exploration arena (91×52.6 cm) into 400 arbitrary bins (20×20 bins), resulting in a bin size of 91/20 cm/bins = 4.55 cm along X and 52.6/20 cm/bins = 2.63 cm along Y). Exploration duration (s) in a given user selected (target zone, T) zone was quantified as the time spent in zone T (*versus* NT, non-target zone) per entry. Body speed over time was computed as the elapsed distance (in cm) and resampled at 10 Hz with a linear interpolation method in MATLAB (non-overlapping 100 ms/window). Data points were excluded if body speed was > 50 cm/s, because we observed that almost all samples fell well under this value (data not shown). The spatial distribution of body speed was computed as a bin-average for each given location and for zone T (*versus* NT) as described above. The animal was considered to be immobile if the body speed was < 1 cm/s and mobile otherwise [5,53]. Body direction over time was computed as the vector angle between two subsequent COM points (see equation 1) and for each given location (10×10 bins), resulting in 91/10 cm/bins = 9.1 cm along X and 52.6/10 = 5.26 cm along Y. Body directions for immobile movements were excluded. A polar frequency histogram of body directions (bin size = 10 deg/bin) was computed as the number of observations normalized by the total number of samples per session.

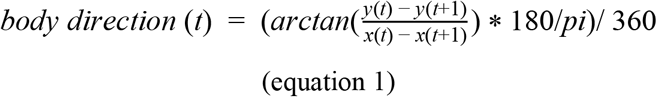

To quantify the relative change in locomotion in T (versus NT), a locomotion modulation index (LMI) was computed as the difference in exploration duration (ED) per entry for T versus NT zone divided by the sum for each sample over time (see equation 2) after data is binned (bin size = 100 ms). Here, the exploration duration was interpolated over time using a linear interpolation method, so that LMI could be computed for each time point. To correct for the difference in surface area of the T and NT zone and avoid a bias for the larger zone in the LMI calculation, we normalized LMI by the surface area. In the current study, the target zone was 45% of the total surface area. The LMI values were corrected accordingly.

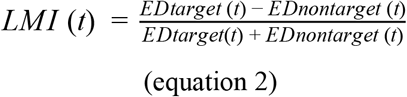

#### 2.5.2 Error analyses

We performed three independent analyses for error estimation. First, to quantify the noise in spatiotemporal motion profiling, we calculated the roboball’s trajectory as described above. The sampled X,Y coordinates from the second sensor device were then rotated 180 degree clockwise around its origin (X,Y = 0,0), to align the world-centric coordinates of detected motion across the two sensor (IR touch) frames (see section 2.2). To remove noise caused by scattering light from the other IR touch frame samples (N = 40/143736 points for dev1; N = 6/32769 points for dev2) the samples that laid outside the robot’s radius (3.81 cm) were removed. The motion trajectories were then resampled at 200 Hz and interpolated by fitting a spline across non-overlapping 50 s windows using the fit function in MATLAB (smoothing parameter = 0.8). Data points were excluded if the traversed speed was > 50 cm/s.

The two IR touch frames were connected to two separate computers that were not synchronized. Therefore the time series from the two sensors were aligned manually. Subsequently, the error in the estimated robot position was computed as the pairwise difference between the two X,Y estimates. Finally, the data was presented as Z scores (see equation 3a-b) to quantify the variance across the two sensors.

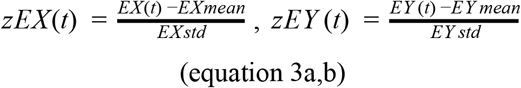

Second, to determine the temporal variation in the spatial estimation, we digitized the location of a stationary object (an orange) using a single IR sensor frame (session duration = 10 min). Temporal variation in spatial estimates was computed by taking the derivative of the COM (Supplemental Figure 3). Finally, to compare the performance in tracking the animal X,Y position between our system with a customized video tracking system as described above, we calculated the difference of the z-scored X and Y samples between video versus sensor tracking (same as in equation 3a,b) and averaged the z-scores in X and Y to obtain a single error value for all sample points over time.

#### 2.5.3 Statistical analyses

To assess place avoidance behaviour, the mobility and body speed were compared when the animal was in zone T versus NT across different tone stimulus conditions (0, 39, 49, 59, random dB) after testing for normality, using the Kolmogorov-Smirnov test. As both variables failed to pass the normality test, the non-parametric Mann-Whitney-U test was used to compare the median exploration duration per entry between zone T and NT for each condition as well as the median body speed. To reduce the risk of type I errors due to multiple testing, p-values were adjusted with a Bonferroni correction, with a critical p-value of 0.05 before correction. To evaluate if there was a bias in the animal’s spatial trajectory, the histogram of heading direction samples was tested with a goodness-of-fit chi-square test (bin size = 10 deg, number of bins = 36, critical p-value = 0.05). A post hoc Pearson’s chi-square test was used for each bin versus the sum of other bins (ratio 1:35) to determine if the observed ratio exceeded the expected ratio (number of samples/number of bins, p-values adjusted with Bonferroni correction).

## 3. Results

### 3.1. Stationary and mobile inanimate object tracking in open field

To evaluate the tracking performance of PolyTouch, we performed three independent analyses. First, we quantified the noise in spatiotemporal estimates by tracking a roboball using two IR sensor frames placed on top of each other (session duration = 15 min, Figure 2A-C) and acquired data simultaneously (see Materials and Methods for details). Noise in the raw samples was quantified as the pairwise difference of z-scores for each axes across the two sensors (Figure 2D). The mean error estimation in the X position was 0.8 cm (± standard deviation, s.d. = 3.6 cm) and 1.0 cm in the y position (± s.d. = 3.5 cm). The proportion of samples with a significant (critical z-score = 1.96 s.d.) error estimation in the body position was 2622/169339 (1.6%) in X, and 502/169339 (0.3%) in Y. After interpolation, the mean error in X reduced to 0.3 cm (± s.d. = 3.5 cm) and 0.2 cm in Y (± s.d. = 3.4 cm). The proportion of samples with a significant error in X was 2407/169339 (1.4%), 543/169339 (0.3%) in Y. These findings show that 98.6% of contacts detected spatiotemporally well matched across sensors.

The temporal variation in the position estimate was quantified independently by tracking a stationary object (an orange) using a single IR sensor frame (session duration = 10 min) with an average sampling rate of ~74 Hz. The temporal variability in COM estimates was zero pixels (Supplemental Figure 3). Finally, to compare and validate PolyTouch’s tracking performance with an existing standard method, we quantified the location estimate of a freely moving mouse by PolyTouch (online tracking at 65.6 Hz; 156.2 Hz in the current version) with a custom video tracking system (offline tracking at 30 fps, see Section 2.3). This revealed a similar performance in tracking the animal position (Figure 2E-F). Although absolute differences were as large as <6 cm (Figure 2F), only a small portion of samples (~0.8%) significantly deviated (|∆Z| > 1.96; Figure 2F). Taken together, our findings indicate that PolyTouch estimated the position of a stationary object with single-pixel precision (Supplemental Figure 3) and performed motion tracking of a moving object with low but variable spatial noise (Figure 2D). Increased error in COM estimate during object motion is primarily because the two sensors sample the same animal across two different planes. While one sensor is placed on the ground where the animal explores, hence sensitive to animal droppings and capture a larger variety of tail movements, the other sensor is elevated off the surface. Because we calculate the error between the sensors after COM calculation, the variance in the estimate is sensitive to the different planes digitized by the two sensors.

**Figure 2.**
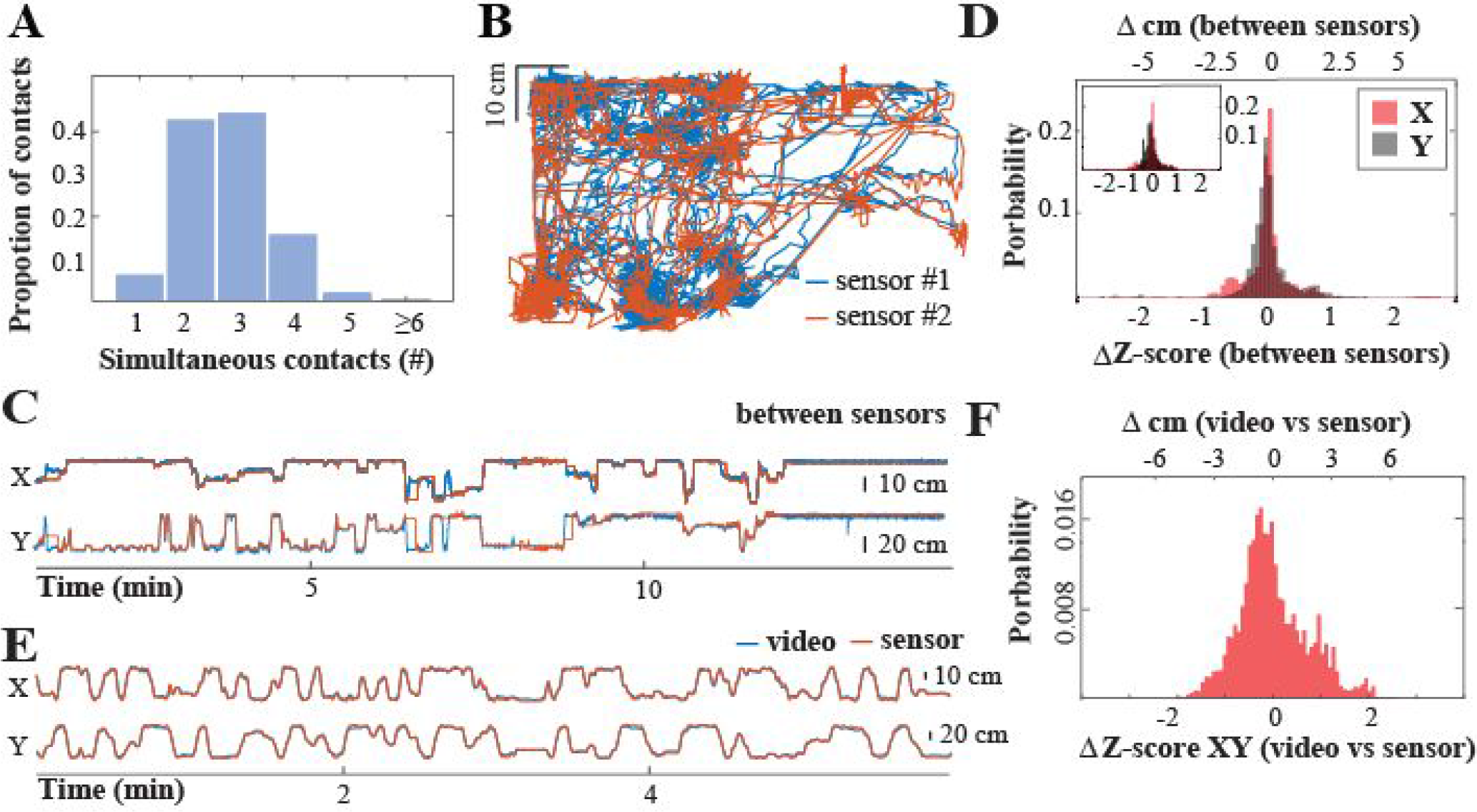
The quantitative assessment of PolyTouch performance for motion tracking. **(A)** Histogram of touch events. Majority of the samples involved 2 (38.8%), 3 (40.2%) or 4 (14.1%) contacts (total number of contacts = 55207). Note that a small fraction of contact events (# contacts > 4) may include the animal’s tail or head, in addition to the four limbs. **(B)** Motion trajectories, quantified as the temporal change in center of mass of all contact points, by two sensors (sensor #1 in blue, sensor #2 in orange). **(C)** The relative distance of the robot from the start position (in cm) along the x-axis (left panel) and y-axis (right panel) over time across the two sensors. **(D)** The Z-scored and absolute cm error difference (see Section 2.5.3) in the COM X (red shaded bars) and Y (grey) across sensors with raw data (small upper panel) and spline fitted data for noise suppression (main panel). Mean ± standard deviation from the mean error before interpolation: −0.8 cm in X (± 3.6 cm) and 1.0 cm in Y (± 3.5 cm); Mean error after interpolation: −0.3 cm in X (± 3.5 cm) and −0.2 cm in Y (± 3.4 cm). **(E)** As in C but the comparison is between the trajectories digitized by PolyTouch (red) and from video recordings (blue). **(F)** Comparison of the animal location as detected by PolyTouch or in video recordings (see Section 2.5.3).

### 3.2. Tracking animal navigation in open field

Next, to exemplify the utility of PolyTouch for closed-loop control of animal navigation, we provided tone feedback every time the animal entered a user-defined virtual target zone. The training consisted of five sessions (t = 5-20 min/session) and is based on a modified open field task [53]. The first three sessions were delivered on the same day (average intersession interval = 7 min) and the subsequent two sessions 5 and 10 days, respectively, after the first experimental day.

Mice could freely explore the arena during the first session (t = 5 min; habituation session) without any tone feedback provided. During the subsequent three sessions, animal entry to a user-defined virtual target zone triggered a discrete 150 Hz tone at 39, 49, or 59 dB (t = 15-20 min/session). During the final session 10s tone sweeps (3x per stimulus condition) were pseudo-randomly presented to decouple the stimulus presentation from the animal’s location (t = 15-20 min).

Locomotion tracking showed that animals remained mobile, i.e. body speed > 1 cm/s, on average 81% of the first session (Figure 3A-B, 3D-E). Animals tended to walk faster along the walls and center part of the arena (Figure 3F-G), and spent the most time at the four corners (Figure 3C). This stereotypical thigmotaxic navigation discontinued in the subsequent sessions as the animals habituated to the environment and became increasingly less mobile.

**Figure 3.**
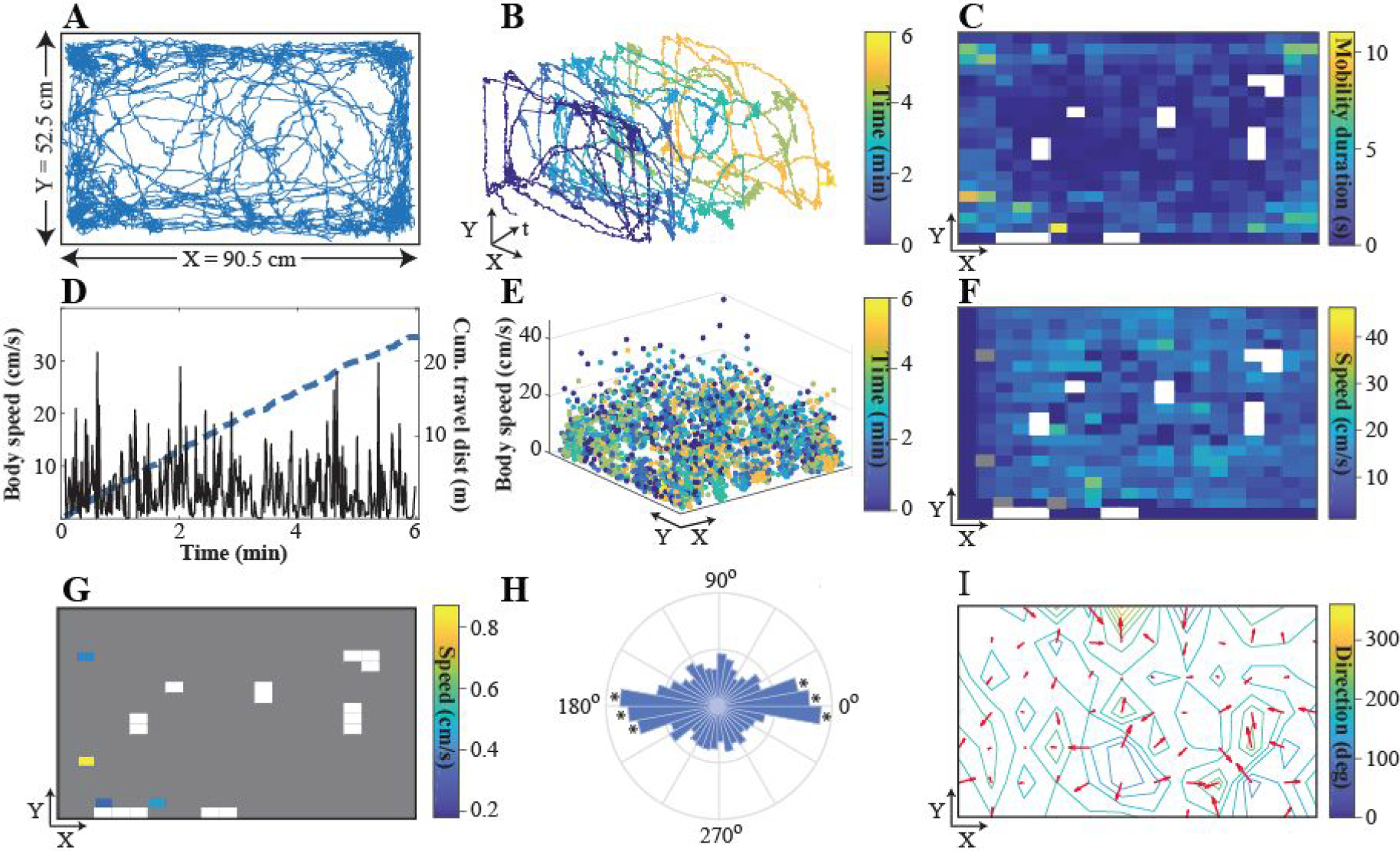
Open field exploration. **(A)** Example trajectory of one mouse (id#2) in the open field during the habituation session (no auditory feedback, session duration = 6 min). The body position is estimated as the center-of-mass (COM) of simultaneous contact points. **(B)** The same trajectory as in (A), but resolved over time. **(C)** Mobility duration (in s; bin size X = 4.55cm, bin size Y = 2.63 cm). **(D**) Time-resolved body speed (black) and cumulative distance traveled (blue) throughout the session. **(E)** Spatial mapping of body speed within arena across the session (spatial resolution = 0.050 cm/pixel). **(F)** Spatial distribution of body speed during animal mobility (bin size X = 4.55, bin size Y = 2.63 cm; white bins indicate never visited positions, grey bins indicate animal mobility was <1 cm/s). **(G)** Same as in (F), but when the animal was immobile (body speed < 1 cm/s; grey bins indicate the animal was mobile). **(H)** Polar histogram of the body direction (bin size = 10 deg) when the animal was moving (* indicates that bin significantly exceeded the expected uniform distribution with p < 0.05, Pearson’s chi-square test). **(I)** Bin-averaged body direction (in deg) and direction gradient (∂F/∂x) from 0 deg (blue) to 360 (yellow) for each given location (bin size X = 9.10 cm, bin size Y = 5.26 cm).

To quantify whether the spatial (thigmotaxic) bias in exploration is accompanied by a motor bias, we computed the body direction (Figure 3H) and the trajectory of animals (Figure 3I). The results showed that once the navigation is initiated, animals either preserved their body orientation, continuing their forward motion, or reversed their trajectory to revisit the path they took (Figure 3H). This pattern of exploration was not spatially constrained to any given portion of the arena (Figure 3I). These results suggest during open field exploration animal behavior is intrinsically driven rather than being extrinsically instructed in this simple arena [53–55].

### 3.3. Place avoidance training with real-time positional feedback

Next, to establish an instructive control over animal navigation, we provided real-time contextual feedback. Animals rapidly learned to avoid the target zone during sessions where entry to a user-defined target zone triggered a moderate intensity (~59 dB) tone (Figure 4). In this close-loop positional (discrete) feedback training, animals first explored the entire arena (Figure 4B) with thigmotaxis (Figure 4B,C) before completely avoiding the target zone after ~7 min exposure (Figure 4B).

**Figure 4.**
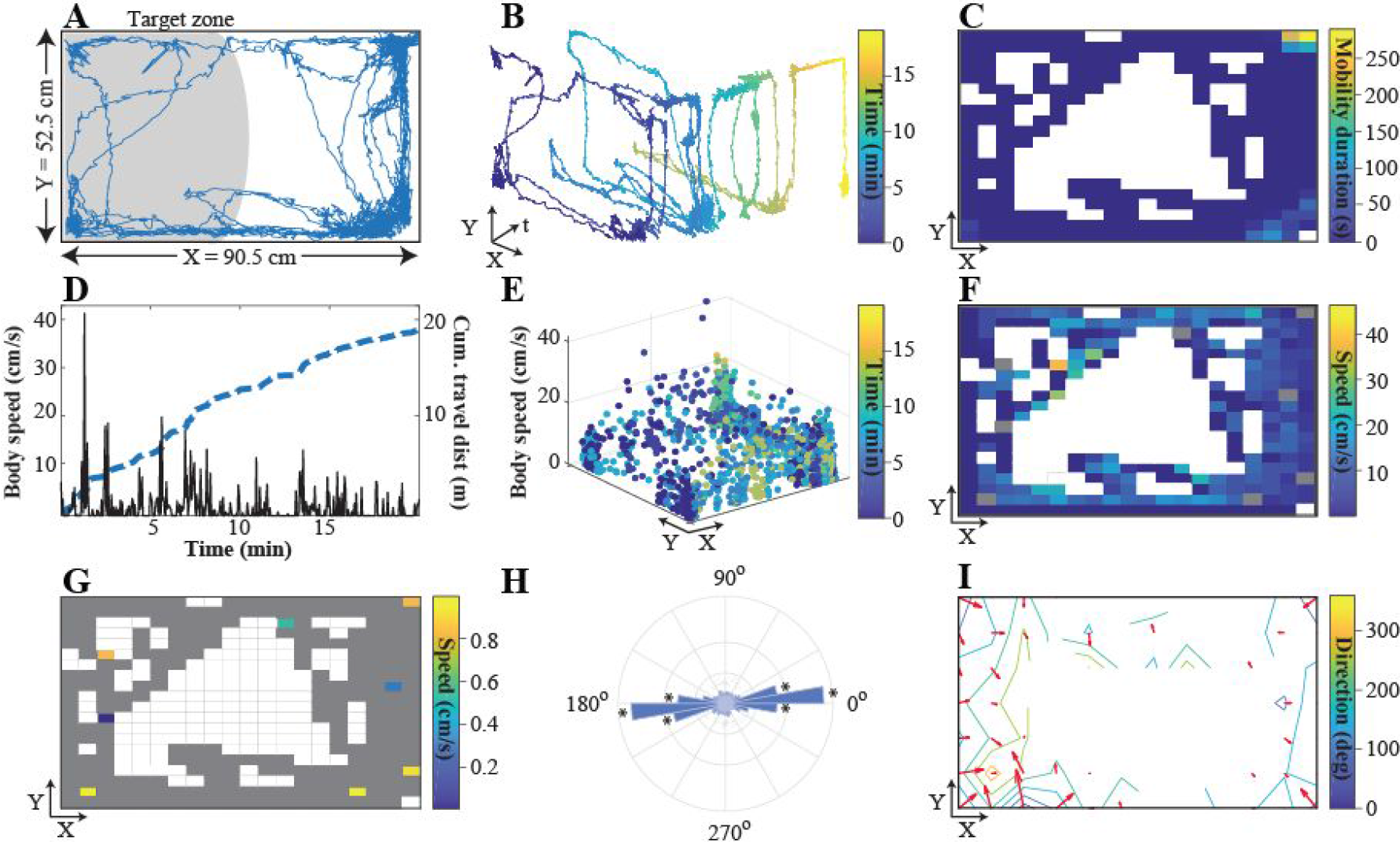
Place conditioning (spatial avoidance training) with close-loop positional (discrete) feedback using virtual targets. **(A)** An example trajectory of one mouse (id#2, same as in Figure 3) when a discrete 59 dB tone feedback was delivered every time animal visited a user selected target zone, shaded in grey (session duration = 20 min). **(B)** The same trajectory as in (A), but displayed over time. **(C)** Mobility duration (bin size X = 4.55cm, bin size Y = 2.63 cm). **(D)** Time-resolved body speed and cumulative distance traveled throughout the session. **(E)** Spatial mapping of body speed within the arena throughout the session (spatial resolution = 0.050 cm). The color code represents time elapsed since the session onset. **(F)** Bin-averaged body speed when the animal was moving (body speed > 1 cm/s) for each given location (bin size X = 4.55 cm, bin size Y = 2.63 cm; white bins indicate never visited positions, grey bins indicate that the animal was immobile). **(G)**Same as in (F), but when the animal was immobile (body speed < 1 cm/s; grey bins indicate the animal was mobile). **(H)** Polar histogram of the body direction (bin size = 10 deg) when the animal was moving (* indicates that bin significantly exceeded the expected uniform distribution with p < 0.05, Pearson’s chi-square test). **(I)** Bin-averaged body direction (in deg) and direction gradient (∂F/∂x) from 0 deg (blue) to 360 (yellow) for each given location (bin size X = 9.10 cm, bin size Y = 5.26cm; white bins indicate never visited locations).

The positional feedback significantly reduced the overall speed of navigation (median ± 1st-3rd interquartile range (IQR) of body speed with and without tone feedback: 0.1 cm/s ± 0-1.8 cm/s vs 3.6 cm/s ± 1.1-8.9 cm/s, Mann-Whitney-U test, p < 0.05) except when exploring the target zone, which was increased during target zone exploration (median ± 1st-3rd IQR of body speed in the T and NT zones: 3.5 cm/s ± 1.0-8.4 cm/s vs 0.2 cm/s ± 0-2.4 cm/s, Mann-Whitney-U test, p < 0.05). Notably, the observed bias in the animal’s trajectory for forward and reverse-traversed motion in the baseline condition (Figure 4H) seemed to be reinforced in close-loop conditions (Figure 4H), suggesting that discrete positional feedback might promote motion along a course.

To quantify the temporal evolution of the avoidance behavior, we calculated a locomotion modulation index (see Section 2.5.1). The LMI index is a normalized measure of explorative behavior that could take positive or negative values. Positive values indicate that animals spend more time (Figure 5A) in the target (T) zone (Figure 5B), while negative values are measures for non-target (NT) zone exploration. The results show that in the absence of auditory feedback (Figure 5, first column) the animals explored both T and NT throughout the entire session (see Figure 6 for group analysis), without a bias towards the exploration of either zone. Low-intensity auditory (39 dB) feedback did not change the exploration duration or the speed of navigation (Figure 5, second column). At higher intensities (49 and 59 dB) mobility was directed towards the non-target zone (Figure 5, third and fourth columns) as animals avoided the T zone only after one entry. After task acquisition, the avoidance behavior was generalized. Animals avoided entry and exploration of the T zone even if the auditory stimulus was not coupled to the T zone entry, rather 10 s tone sweeps were delivered randomly during exploration (Figure 5, last column).

**Figure 5.**
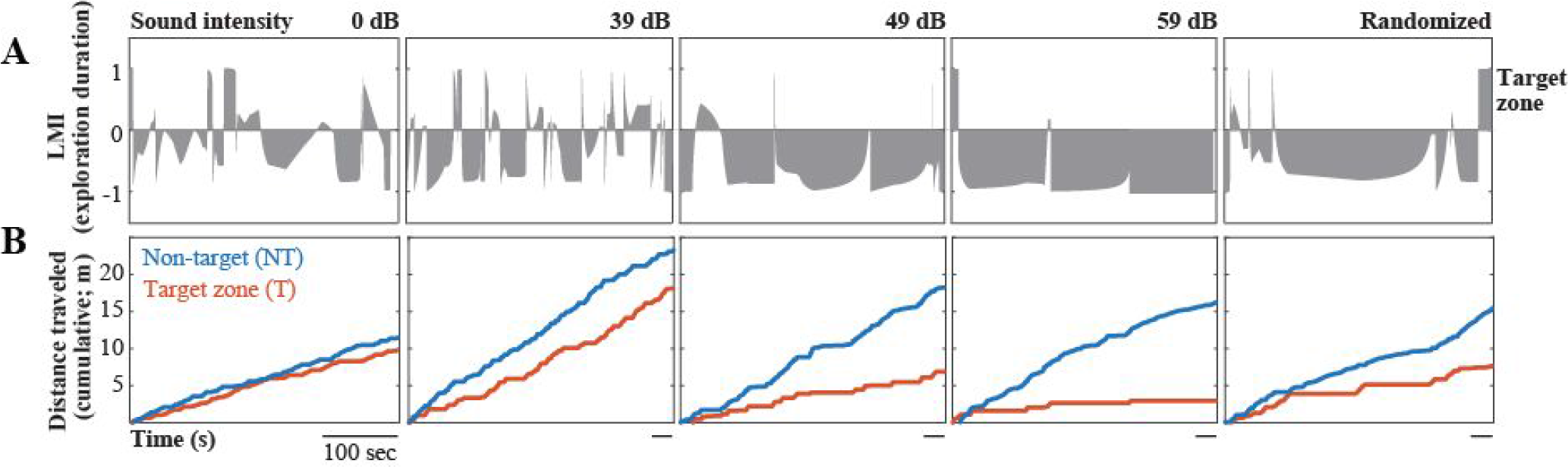
Temporal evolution of the place conditioning. Open field navigation was quantified in the absence of auditory feedback (baseline, 0 dB), or during auditory feedback for one mouse (id#2). The feedback was provided either in a close-loop whenever the animal entered the virtual target zone, using three different tone intensities (39-49-59 dB), or as open-loop feedback, presented pseudorandomly over time (3x 39, 49, 59 dB 10 s tone), independent from the animal’s visit of the virtual target. **(A)** Locomotion modulation index (LMI) for the duration of time spent in the T and NT zones. LMI = ED_T_ − ED_NT_/ED_T_ +ED_NT_, (see Section 2.5.1). Positive values denote preferential exploration of the T zone, negative values NT zone. **(B)** Cumulative distance traveled across each session. Data from a single representative animal.

Group analyses showed that animal mobility was modulated by both close- and open-loop auditory feedback (Figure 6). During close-loop feedback, when auditory stimulus was delivered at higher intensities, the animals explored the T zone less, avoiding the virtual target (Figure 6A; p<0.05, Mann-Whitney-U test), and moved faster (Figure 6B; p<0.05, Mann-Whitney-U test). Faster mobility was also observed during open-loop feedback (Figure 6B; p<0.05, Mann-Whitney-U test). Taken together results outlined in Figures 4–6 argue that place conditioning can be successfully induced during spatial avoidance training using virtual target zones as PolyTouch provides context/location specific and independent feedback. The place avoidance is rapidly induced in a single session and can be generalized across stimulus intensities.

**Figure 6.**
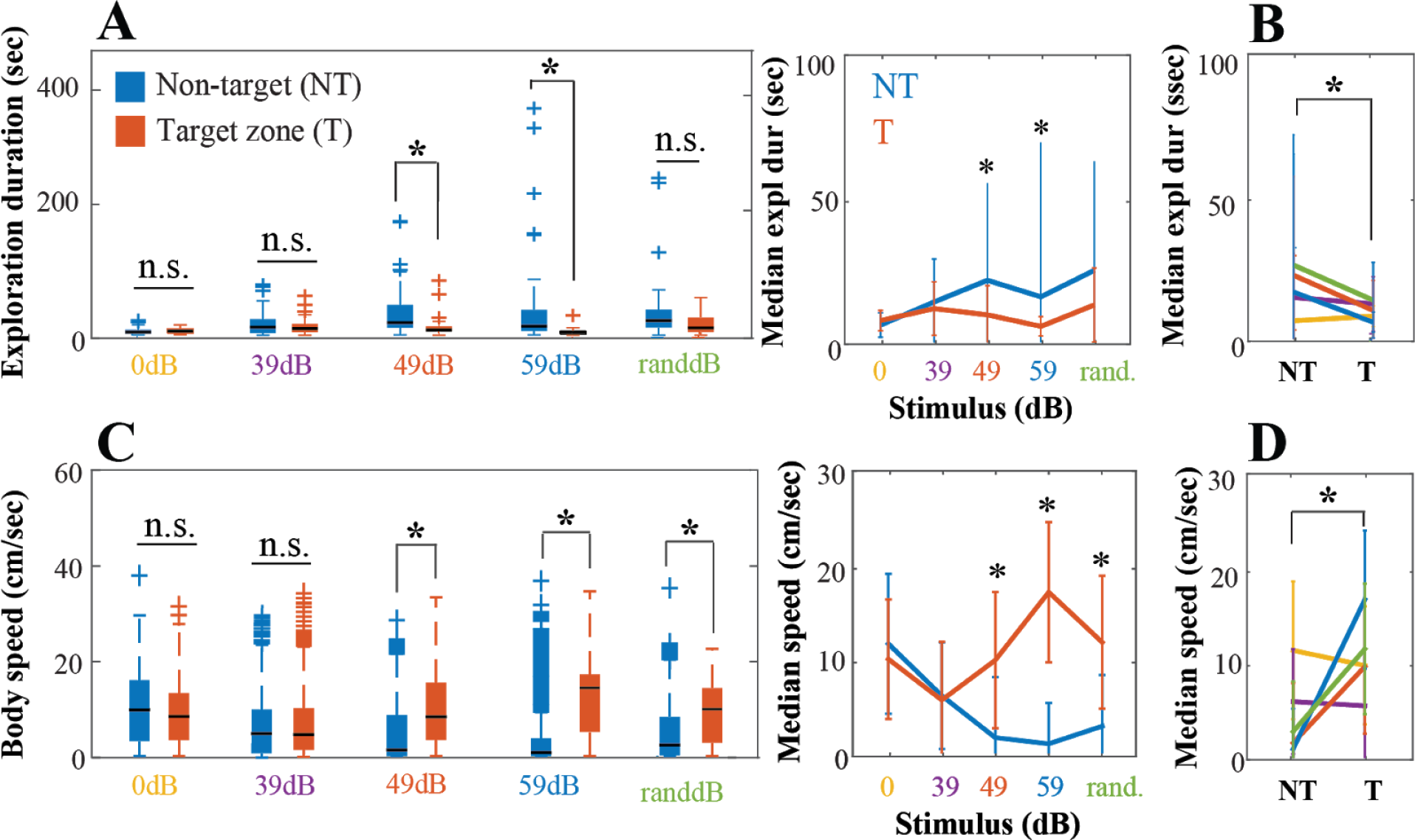
Comparison of animal mobility across stimulus conditions. (A) Exploration duration (ED) pooled across two mice (id#1 and 2) in the randomly assigned target (T, orange) versus non-target (NT, blue) zone baseline, 39, 49, 59 and random dB tone conditions. Left panel: boxplot with median ED (black line), 1st and 3rd quartile outliers (lower and upper lines), and outliers (cross symbols). The median ED was not significantly different between the T and NT zone for the baseline (ED_T_ = 7.9 s, 1st-3rd interquartile range, IQR_T_ = 4.5-11.8 s; ED_NT_ = 6.3 s, IQR_NT_ = 3.4-10.2 s; p = 0.98), 39 dB (ED_T_ = 12.2 s, IQR_T_ = 5.9-19.7 s; ED_NT_ = 14.6 s, IQR_NT_ = 4.5-28.2 s; p = 1) and random tone condition (ED_T_ = 13.6 s, IQR_T_ = 6.3-31.3 s; ED_NT_ = 26.1 s, IQR_NT_ = 14.0-44.5 s; p = 0.066), but was significantly lower in the T zone compared to NT zone for the 49 dB (ED_T_ = 10.0 s, IQR_T_ =7.8-16.4 s; ED_NT_ = 22.4 s, IQR_NT_ = 13.7-52.7 s; p = 7.3E-04) and 59 dB condition (ED_T_ = 5.8 s, IQR_T_ = 3.0-7.7 s; ED_NT_ = 16.5 s, IQR_NT_ = 8.3-44.2 s; p = 2.2E-06). (B) The median ED pooled across tone conditions was significantly higher lower for zone T compared to NT (median_T_ = 9.0 s, interquartile range (IQR_T_) = 5.2-16.8 s; median_NT_ = 14.7 s, IQR_NT_ = 6.3-31.3 s, p = 2.4E-05). (C) Same as in A but for body speed (BS, cm/s). The median BS was not significantly different between the T compared to NT zone for the baseline (BS_T_ = 9.5 cm/s, IQR_T_ = 3.8-15.0cm/s; BS_NT_ = 11.0cm/s, IQR_NT_ = 3.6-18.1cm/s; p = 0.09) and 39 dB tone condition (BS_T_ = 5.3 cm/s, IQR_T_ = 1.7-11.6 cm/s; BS_NT_ = 5.7 cm/s, IQR_NT_ = 1.1-11.5 cm/s; p = 0.9). In contrast, the median BS was significantly higher for the T zone for the 49 dB (BS_T_ =9.4 cm/s, IQR_T_ = 3.9-19.5 cm/s; BS_NT_ = 1.5 cm/s, IQR_NT_ = 0-9.8 cm/s; p = 7.6E-100), 59 dB (BS_T_ = 16.3 cm/s, IQR_T_ = 5.8-19.4 cm/s; BS_NT_ = 0.8 cm/s, IQR_NT_ = 0-4.2 cm/s; p = 4.8E-233), and random tone condition (BS_T_ = 11.2 cm/s, IQR_T_ = 3.2-16.2 cm/s; BS_NT_ = 2.6 cm/s, IQR_NT_ = 0.2-9.4 cm/s; p = 3.6E-122). (D) The median BS pooled across tone conditions was significantly higher lower for zone T vs NT (median_T_ = 9.9 cm/s, IQR_T_ = 2.8-17.0 cm/s; median_NT_ = 2.5 cm/s, IQR_NT_ = 0.1-9.8 cm/s; p < 0.0001). A Mann-Whitney-U test was used for all multiple comparisons (see Section 2.5.3 for the details on statistical method selection). * denotes p<0.01.

### 3.4. Continuous positional feedback during open field navigation

Discrete two-state feedback provides animals binary information, e.g. whether or not they are in a target zone as implemented in Section 3.3. PolyTouch can also be used to provide higher dimensional and continuous feedback. We have implemented such a training protocol by continuously providing auditory feedback regarding an animal’s relevant location to a user selected virtual target zone (Figure 7). During the habituation session, a mouse explored the open field (t = 5 min) without any auditory feedback. In the next two sessions, the subject received continuous tone that increased (or decreased) in frequency (150:150:750 Hz or 750:-150:150 Hz) as it approached the target (t = 10 min/session; average inter-session interval = 5-10 min).

To determine how the animal’s exploration behavior changed as a function of the distance to the target zone, we quantified the exploration duration (in s), body speed (in cm/s), and body direction (in deg) over time and for each zone in the open field. The walked trajectory revealed that the animal actively explored the arena during the baseline session (data not shown), with a proportion of time spent in a mobile state of 0.90 (body speed > 1 cm/s). The animal spent most time in zone 2 (exploration duration, ED = 104.2/298.9 s = 34.9%) and zone 5 (ED = 82.5/298.9 s = 27.6%; Figure 7J, Supplemental Figure 8A), spending the most time at the four corners.

The body speed varied across zones, but no obvious differences were observed (Supplemental Figure 8B). Similarly, the animal remained mobile in the subsequent session and spent most time in zone 2 (ED = 242.3/702 s = 34.5%) and zone 5 (ED = 250.3 /702 s = 35.7%; Figure 7K, Supplemental Figure 8A). Again, no obvious differences were observed in the body speed of the animal across zones (Supplemental Figure 8B). This may indicate that exploration behavior under closed-loop conditions was similar to baseline conditions. Indeed, the heading direction of the animal’s trajectory revealed similar frequency distributions and was non-uniformly distributed (p < 0.05, Pearson’s chi-square test).

In contrast, after the first session of continuous positional feedback training, the animal systematically alternated between mobile and immobile periods (proportion immobile = 0.56, body speed < 1 cm/s; Figure 7A-B, D). It spent the most time exploring the zone 5 (ED = 369 /710 s = 51.9%; Figure 7C, Supplemental Figure 8A), where he moved the slowest (Supplemental Figure 8B). The frequency distribution of the animal’s heading direction was non-uniformly distributed (p < 0.05, goodness-of-fit chi-square test, Figure 7H). Given that the lowest frequency tone was presented in zone 5, our findings may indicate that the animal preferred to stay in the area that was coupled with the lowest sound intensity feedback.

**Figure 7.**
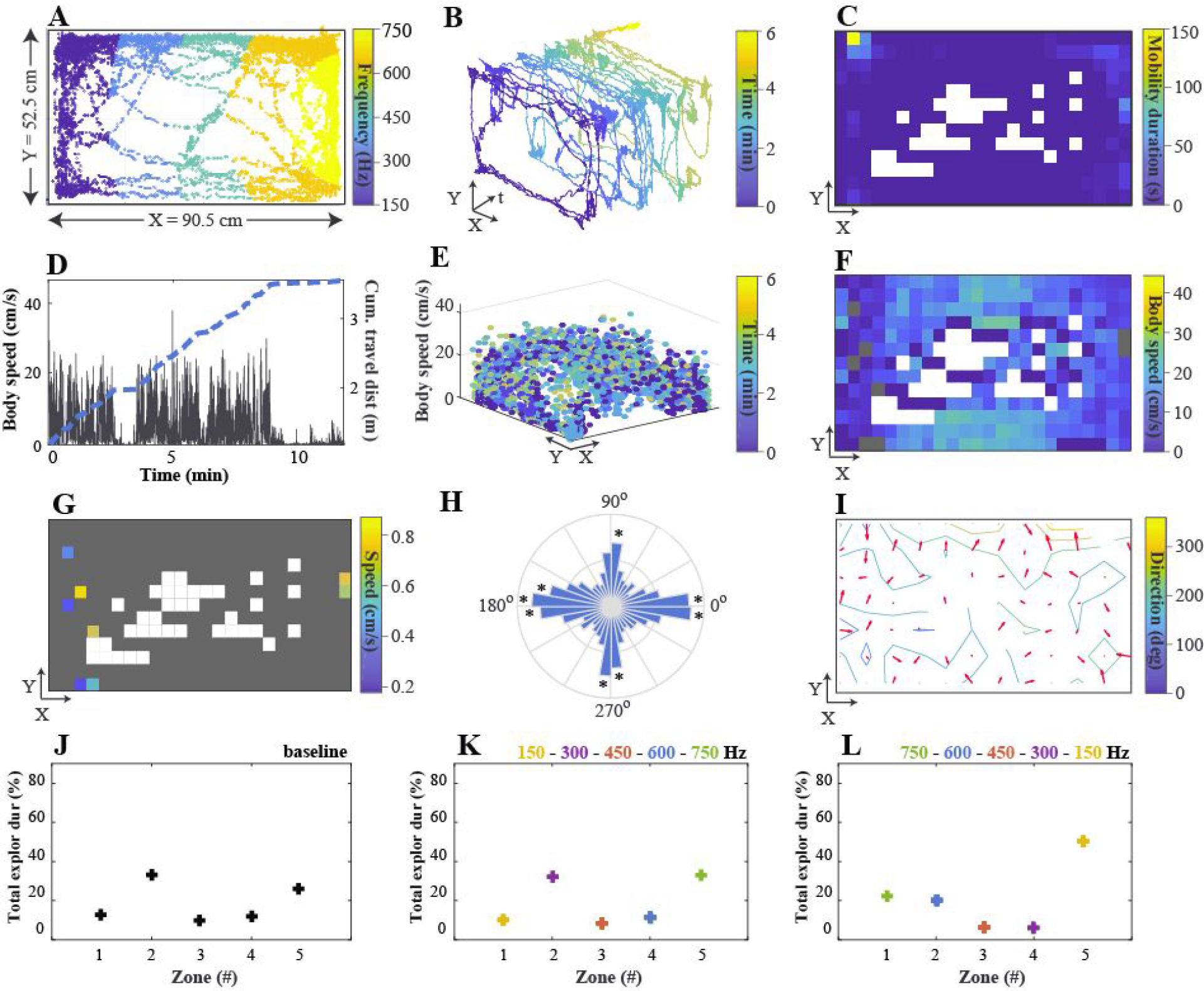
Locomotion activity in an open field with close-loop auditory feedback. **(A)** Example trajectory of one mouse (id#3) in the open field where a continuous sine tone was given with a frequency that scaled with the distance of the animal to five different virtual target zones (session 3, session duration = 10 min). Each dot represents the center-of-mass (COM) body position. **(B)** Same as in (A), but over time (t, in min). **(C)** Bin-sum mobility duration (MD, in s) for each given location in the open field (bin size = 3.93 cm/pixel; white bins indicate never visited positions). **(D)** Body speed (in cm/s) over time (in min) after applying a linear interpolation function (non-overlapping 3 s/bin windows). **(E)** Body speed (in cm/s) as a function of the center-of-mass position in the open field throughout the session. **(F)** Bin-averaged body speed when the animal was moving (body speed > 1 cm/s) for each given location (bin size = 3.93 cm/pixel; white bins indicate never visited positions, grey bins indicate that the animal was immobile). **(G)** Same as in (F), but when the animal was immobile (body speed < 1 cm/s; grey bins indicate the animal was mobile). **(H)** Polar histogram of the body direction (in deg, bin size = 10 deg) when the animal was moving. **(I)** Bin-averaged body direction (in deg) and direction gradient (∂F/∂x) from 0 deg (blue) to 360 (yellow) for each given location (bin size = 7.54 cm/pixel; white bins indicate never visited locations). **(J)** Total MD as a percentage of the session duration (in %) in each zone 1 to 5 for the baseline session (no feedback): 13.8%, 34.9%, 11.2%, 12.6% and 27.6%, respectively **(K)** Same as in (J), but for the closed-loop condition where tone feedback decreased in frequency as the animal approached zone 5. Total MD as a percentage of session duration for zone 1 to 5: 10.1%, 34.5%, 8.6%, 11.2% and 35.7%, respectively **(L)** Same as (J), but where tone feedback increased in frequency as the animal approached zone 5. Total MD as a percentage of session duration for zone 1 to 5: 21.1%, 19.1%, 3.8%, 4.0% and 52.0%, respectively.

## 4. Discussion

PolyTouch is a novel, open source, markerless (i.e. does not require body part of animals to be tagged) tracking software with integrated feedback delivery capabilities. It continuously tracks animal locomotion while providing control signals to connected devices for closed-loop control (Figure 1A-B). The main features of the software are (1) simultaneous tracking of multiple body parts using any IR sensor frame, connected to the data acquisition computer via the Universal USB Touchscreen Controller driver, with an average sampling rate of 156.2 Hz (in the current version of the software) and fine spatial resolution of 0.050 cm as deployed in this study, (2) rapid millisecond feedback based context or animal position with an average feedback latency of ~5.7 ms, (3) low but variable noise (temporal: ~0%; spatial: <1.7% at cm pixel resolution) reconstruction of navigation over extended periods of time and (4) continuous update of the stored behavioral data with minimal memory load on the host computer resources.

PolyTouch provides users real-time visual feedback by displaying touch locations, center-of-mass (COM) body position, body speed, traveled distance, behavioral state (mobile, immobile), and distance to any user-defined virtual zone. It is fully automated and flexible, as the user can access various locomotion variables during and after data acquisition from a lightweight output file (1 min ~ 79 KB). This provides means to evaluate the occurrence, duration and timing of behavioral sequences in a standardized manner and allows for synchronization with other experimental modules, e.g via provided MATLAB scripts for real-time data visualization, feedback control, integration with electrophysiological recordings. The recording arena can take any size, constrained only with the dimensions of the sensor frame, and could be extended to the third dimension using multiple IR sensor frames simultaneously. The open field arena can be extended with transparent walls and tunnels to create a more complex and enriched environment, which not only widens the variety of exploration behaviors that can be studied but also benefits the animal welfare [56].

We implemented the closed-loop system in two exemplary place awareness paradigms. In the first paradigm, a discrete tone was presented whenever the animal entered a user-defined target zone. In this context, animals spent significantly less time and faster in the target compared to non-target zone, but only if the tone intensity was sufficiently high (Figure 6A-B), indicating that place avoidance behavior emerged if the tone feedback was aversive. The tendency to avoid the target zone (which was randomized across sessions) increased with the exposure time to auditory feedback within the span of a single session (Figure 5A, 5C). This result argues that animals actively switched their place preference based on the sensory feedback within ~20 min as a result of rapid spatial learning [57,58]. In the second paradigm, continuous feedback about an animals’ relevant distance to the target zone was provided by modulating frequency of the tone, providing real-time positional feedback. After the initial familiarization to the paradigm, the animal spent the most time in the portion of the arena where the lowest frequency stimulus was given, maximizing the distance between the target and itself (Figure 7). Taken together, we show that PolyTouch can be used to provide real-time discrete and continuous sensory (auditory) feedback to bias animal navigation with virtual targets. The virtual targets can be spatially anchored in the arena (as it is implemented in the current study) and take any 2D shape or can be extended to 3D by using multiple sensors to track animals’ elevation. Alternatively, the targets can be coupled to animals’ own exploratory body movements, e.g. change in speed and direction of movement, to immobility, or multiple target zones can be designated to shape animal navigation. Inclusion of objects in the exploration arena and creating associated virtual targets can extend the utility of this approach, as auditory stimulus frequency and intensity could be modulated to create aversive contexts.

### 4.1. Limitations of the PolyTouch

The closed-loop system of PolyTouch consists of a tracking and feedback module that run in parallel (Figure 1B). This design choice ensures that temporal delays introduced by the tracking (~6.7 ms) and feedback modules (~6 ms) do not interfere with one another. The sampling rate and latency of feedback, however, depend on a number of factors. The feedback latency increases from ~5.7 (± 1.8, std) ms to ~12.6 ms (± 3.8, std) if the number of bodies in motion detected is increased to ≤4. The sampling rate was ~74 Hz when an immobile object (single point) was tracked compared to ~65.5 Hz in freely moving mice (up to 6 points tracked/mouse, see Figure 2A for the contact distribution) using the first release of PolyTouch. The new release available for download at Github (https://github.com/DepartmentofNeurophysiology/PolyTouch) significantly improves the sampling rate to 156.2 Hz for animal tracking. The sampling rate remains constant within and across sessions, indicating that memory handling by PolyTouch does not interfere with the tracking performance.

Our sampling error analysis revealed a small (1.7%), not statistically significant, error in animal tracking when two frames were used to track the same animal (Figure 2D). There are two possible explanations for the observed errors: 1) the horizontal alignment of sensor frames in a vertical axis might have resulted in light contamination across the sensors which can be eliminated by optically isolating the frames; 2) the two sensors sampled different portions of the body. The sensor placed on the surface was used to track the position of the limbs, and at times the tail, while the top sensor was positioned to digitize body position. Because micromovement of the limbs or the change of the center of body mass without a change in limb position would be detected only by one sensor, the error across the two sensors could also be attributed to micro motor movements. Finally, the use of two separate computers to acquire the data might have contributed to the sampling error as system utilization in each computer will impact the sampling rate, and indirectly, the precision in spatial localization.

PolyTouch does not currently support simultaneous tracking of multiple unrestrained animals. This is because the center of mass calculation is performed in every time point (“frame”) independently. When multiple animals are in close proximity this algorithm will fail to identify single animals. A Bayesian clustering approach might allow identification of the animal ID based on each animal’s motion trajectory and the previous spatial distributions of individual touch events. Another solution might be to position a camera above the exploration chamber so that a post hoc image-based analysis can be performed to assign the correct animal to touch points. Although the latter solution will limit closed-loop feedback capabilities of PolyTouch to spatial feedback, as single animal behavior will not be classified in real-time for rapid feedback, the solution could be implemented with the current version of the PolyTouch.

The feedback module of PolyTouch uses the audio out jack of the computer as a communication port to deliver control signals to external devices. In this study we provided auditory feedback via connected speakers, however, the control signal could take any shape, e.g. TTL.

### 4.2. Comparison to existing tracking systems

Compared to existing open source software for animal tracking, PolyTouch is the only software that enables rapid (i.e. ~5.7 ms) close-loop feedback for behavior (see Table 1 for a detailed comparison). Integration of a matrix sensor frame with wide light beam emitters (see Materials and Methods) allowed us to capture animal locomotion with a higher spatial resolution in comparison to existing infrared beam tracking systems: 0.050 cm (PolyTouch) versus 1.6 cm [57], 2.54 cm [58], 2.63 cm [59] and >5 cm [23,26,60]. A direct comparison of PolyTouch with a customized video system revealed that the accuracy of PolyTouch is similar to manual offline tracking by an experienced human observer (Figure 2E-F), validating the potential use of PolyTouch in comparison to standard tracking systems. In comparison to video-based systems, one advantage of our system is that automated online tracking by PolyTouch does not require time-consuming image processing steps (e.g. color thresholding, contrasting, contouring). Because image formation does not require the use of an objective, unlike video based animal tracking, spatial resolution of animal tracking by PolyTouch remain homogenous throughout the exploration arena, independent from its size.

In terms of processing speed, modern closed-loop systems trigger feedback within 3-40 ms for neural data [11,13] and 12-50 ms for behavioral data [15,16,42,59,60]. PolyTouch can be used to control neural activity based on animal behavior at a temporal scale faster than synaptic communication along sensory pathways [61–66], and thus could be used to create artificial sensory and motor feedback in the context of rapidly evolving behavioral computations [43,67].

In recent years, many researchers have benefitted from increased availability of open-source platforms, including OpenEphys [11], BONSAI [10], NeuroRighter [68] and RTXI [69]. PolyTouch compliments the functionality of these platforms by providing rapid (within, on average, 5.7 ms) closed-loop feedback based on animal behavior. Although our system supports data streaming of behavior to a multitude of downstream devices, it will (1) require manual coding by the user to add new threads of execution and (2) compromise in speed of processing with each extension due to its dependency on the resource utilization of the data acquisition computer (DAC). The latter issue can be solved by running PolyTouch on a separate DAC.

**Table 1.** A comparison of open source software tools for tracking freely moving animals.

| **Publication (year)** | **Tracking system** | **Tracking principle** | **Main features** | **Temporal resolution** | **Real-time** | **Real-time latency** | **Algorithm flexibility** |
| --- | --- | --- | --- | --- | --- | --- | --- |
| Hamers et al.(2006) | CatWalk | IR-light sensors | High-contrast imaging of footprints with a frustrated total internal reflection (FTIR)-based sensor system coupled to a CCD camera | 50-60 fps | no | N/A | Semi-automated: requires initial manual marking of footprints |
| Machado et al.(2015) | MouseLoco | Videography | Offline 3D markerless tracking of whole-body, limbs, head, tail with automated behavioral classification | 400 fps | no | N/A | Fully automated; posthoc automated classification of behavioral events; Synchronization with physiology data (BONSAI) |
| Mendes et al.(2015) | MouseWalker | IR-light sensors, videography | High-contrast imaging of footprints and whole-body with a frustrated total internal reflection (FTIR)-based sensor system coupled to a high-speed camera | 250 fps | no | N/A | Fully automated |
| Del Rosso et al. (2017) | ratCAVE | Videography | 3D tracking of head position in freely moving rats with close-loop visual feedback in a virtual reality system | 240-360 fps | yes | 15 ms | Fully automated; real-time calibration of head position and rotation to image projector; no training required for VR setup |
| Nashaat et al.(2017) | PIXY | Videography | Online and offline color-based tracking of whole-body, limbs, head and whiskers using a hybrid camera system. Real-time color frames are synchronized with replayed frames obtained at high speed | 150 fps high-speed camera; 25-50 Hz Pixy color camera | yes | 30 ms | Fully automated; Inexpensive; includes wrapper for data processing (PixyMon software) |
| Saxena et al.(2018) | PiCamera | Videography | Tracking of whole-body movements in large arenas (5.5x3m) using a scalable camera system | 30 Hz | no | N/A | Fully automated |
| Lim & Celikel (current study) | PolyTouch | IR-light sensors | Online tracking of body position using a low-level GUI-based multi-touch interface with real-time stimulation. Markerless, inexpensive, light/dark conditions. | 156 Hz | yes | 5.7 (if single body tracked) | Fully automated; Inexpensive; standalone; includes wrapper for MATLAB to automate data acquisition, analysis and visualization |

### 4.3. Future directions

PolyTouch is a flexible software whose functionality can be further improved via extension packages. To support detection of additional behaviors, such as rearing and grooming, the recording range can be extended by positioning multiple sensor frames on top of each other, so that horizontal, as well as vertical movements, are monitored. The user will be able to easily group locomotion data according to the position of the frame since our software saves the device identity for each touch input in the output file. For specific behaviors that require manual observation (e.g. grooming, scratching), a future release will include the possibility for the user to report events by means of a specific key press during data acquisition.

As an extension pack, a robotic command module is currently under development that will allow users to control a robot a Sphero robot (ORBOTIX) in closed-loop based on the animal’s spatial trajectory to study robot-animal interactions [18,70], foraging behavior with predator-prey interactions [19,71,72] and guided spatial learning in complex navigation tasks [73]. We hope this free, open-source, fully automatized tracking software will provide users with an alternative low-cost method to track animal behavior and closed-loop control of brain and behavior.

## Supplementary Figures

**Supplemental Figure 1.**
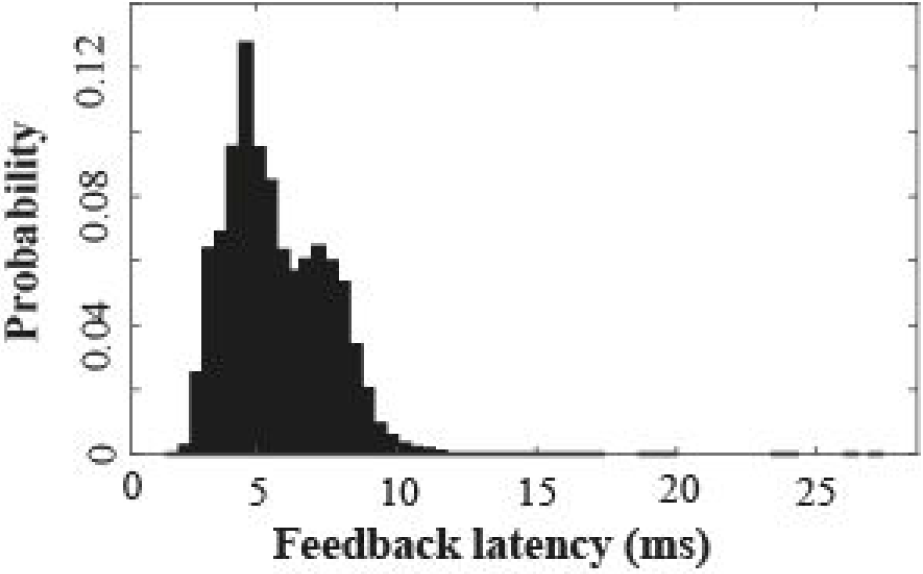
Feedback latency distribution. Normalized histogram of feedback latency (in ms) when a single body in motion is tracked (sampling period = 2 min). Average latency (± standard deviation, std): 5.7 ms (± 1.8 ms).

**Supplemental Figure 2.**
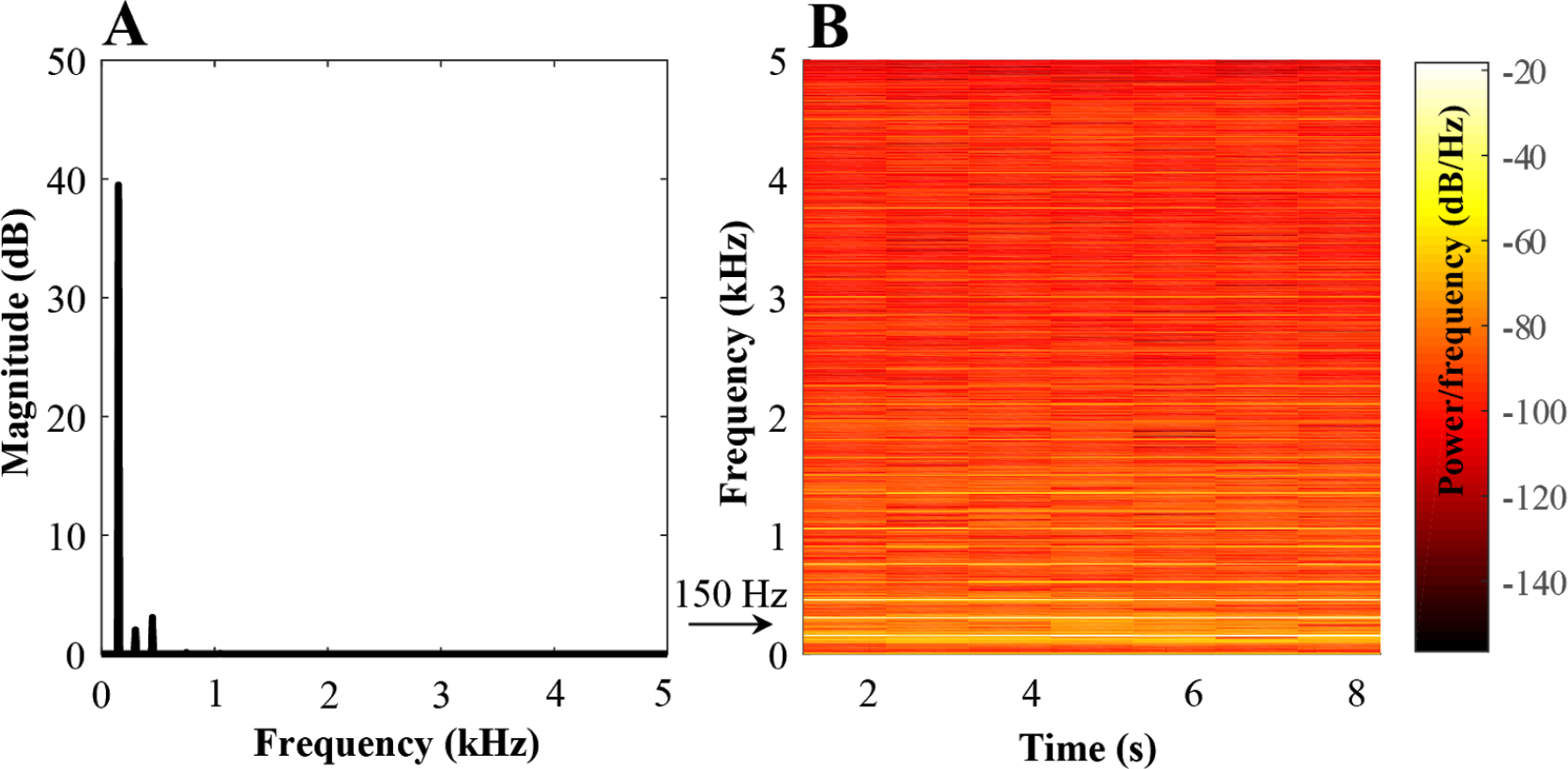
Spectral analysis of sample tone stimulus. **(A)** Single-sided frequency spectrum of a 150 Hz sine wave tone (60 dB) played from two external audio speakers (frequency in kHz, sample duration = 15 s). A fast Fourier transform was computed on the original sound sample and confirmed that the principal frequency was 150 Hz with a magnitude of ~40 (frequency resolution = 10 Hz). The frequency window was limited to 0 to 5 kHz to show relevant frequency peaks. **(B)** Spectrogram of the sound sample described in (A) with frequency (in kHz) over time (in s) using a short-term transform with 50% overlap between adjacent windows (n windows = 8), where colors indicate the power per frequency (in dB/Hz).

**Supplemental Figure 3.**
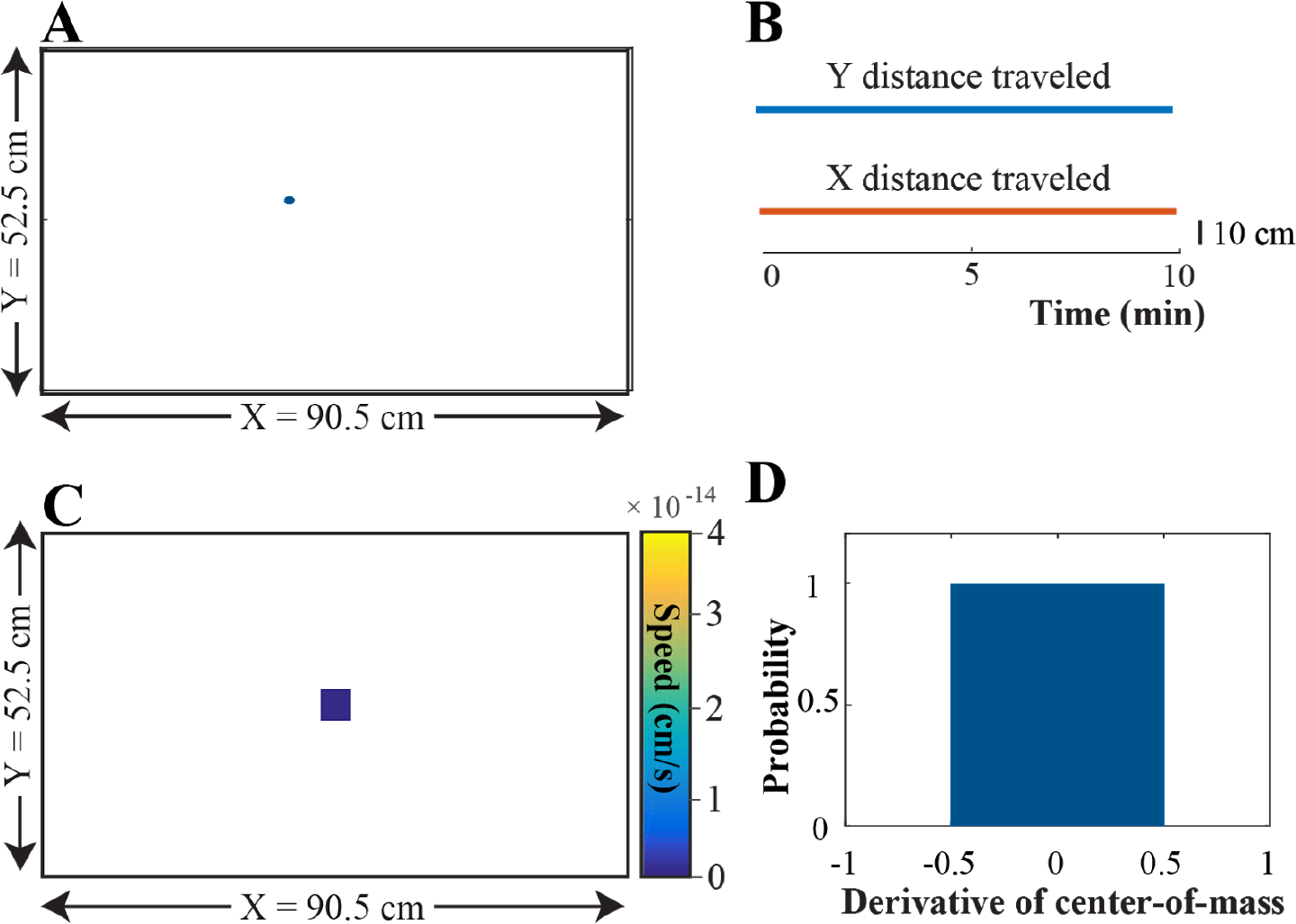
Stationary object tracking in the open field. To evaluate the tracking performance of PolyTouch, we tracked a stationary object (an orange) for error estimation of the spatial X,Y estimates. **(A)** Spatial trajectory of the object in the open field shows that a single X,Y point (X = 788, Y = 557, blue dot) was detected (session duration = 10 min; spatial resolution = 0.050 cm). **(B)** The X,Y coordinates over time (in min) reveal that the object position remained constant over time. **(C)** Bin-averaged body speed (in cm/s) for each given location in the open field (bin size = 3.93 cm/pixels). White bins indicate that the object was not detected in said bin. The average body speed of the immobile object was 0 cm/s. **(D)** Histogram of the derivative of center-of-mass object position reveals zero spatial variation at a single-pixel resolution.

**Supplemental Figure 4.**
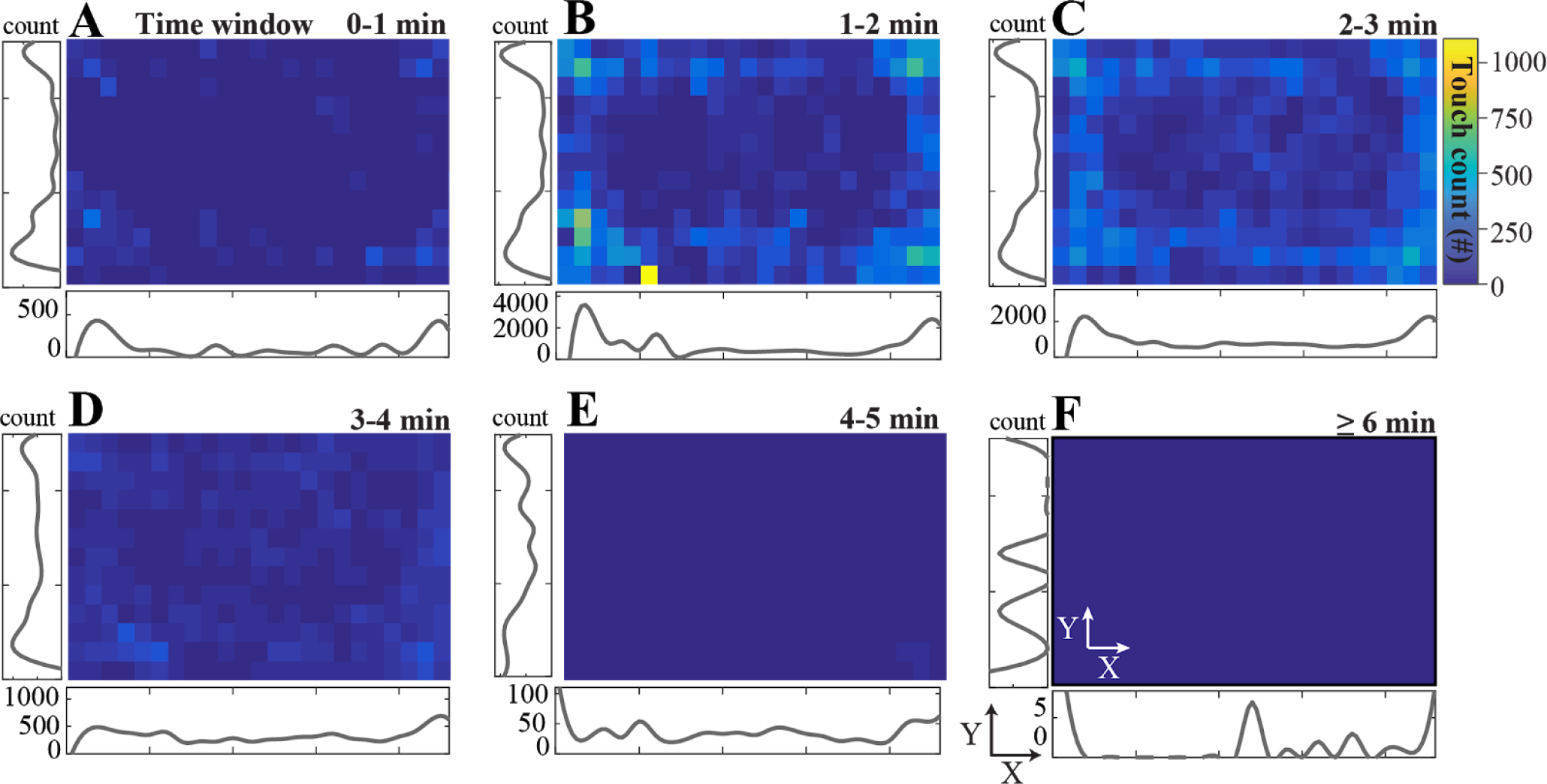
Density of multi-touch events as a function of body position (animal 2). **(A)** Total number of single-touch events (n = 2980) spatially binned as a function of the body position in the open field (bin size = 80×80 pixels) for an example session of animal 2 during the baseline condition (0 dB; session duration = 6 min). Adjacent density plots of touch events (counts) along the x- and y-axis. **(B)** Same as (A) for touch events where 2 coincident touches were detected (n = 21444). **(C)** Same as (A) for touch events where 3 simultaneous touches were detected (n = 22194). **(D)** Same as (A) for touch events where 4 coincident touches were detected (n = 7768). **(E)** Same as (A) for touch events where 5 coincident touches were detected (n = 790). **(F)** Same as (A) for touch events where ≥6 coincident touches were detected (n 6-touches = 24, n 7-touches = 7).

**Supplemental Figure 5.**
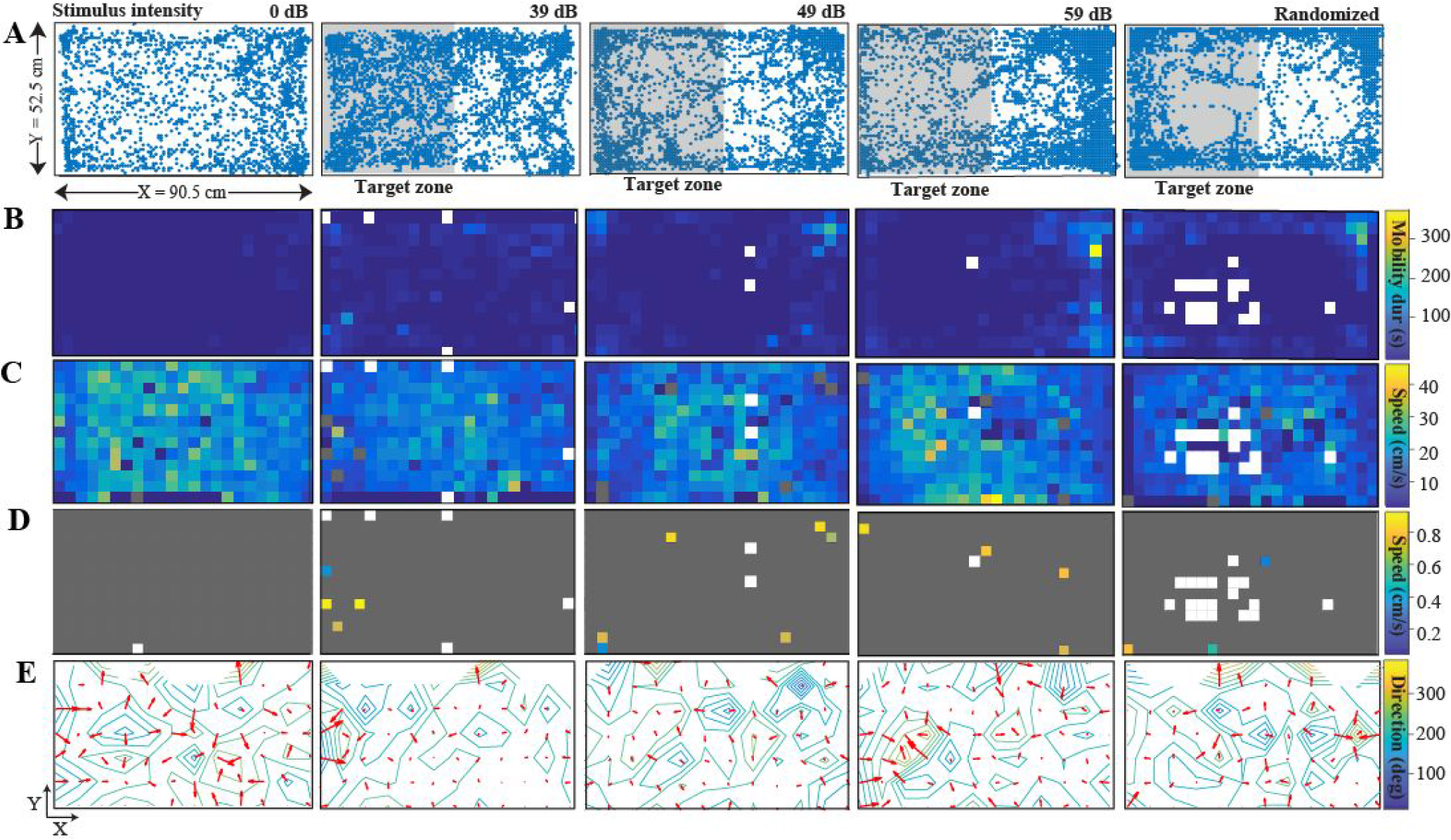
Locomotion activity in open field with discrete closed-loop feedback (animal 1). **(A)** Trajectory of animal 1 in open field arena for 5 different stimulus conditions: no feedback when the animal was in the target (T grey) zone (0 dB, session duration = 6 min, left panel), auditory feedback when the animal was in the T zone (39, 49, 59 dB discrete tone, middle 3 panels), or random feedback given pseudorandomly over time (3x 39, 49, 59 dB 10s tone, right panel). The position is the centre-of-mass of coincident multi-touches (bin size = 0.273 mm/pixel). **(B)** Bin-summed mobility duration (in s) for each given location in the open field (bin size = 80×80 pixels) across stimulus conditions. White bins indicate never visited positions. **(C)** The bin-averaged body speed (BS, in cm/s) for each given location when the animal was mobile (BS > 1 cm/s) across stimulus conditions. White bins indicate never visited places and grey bins indicate that the animal was immobile (BS < 1 cm/s). **(D)** Same as (C), but when the animal was immobile. Grey bins indicate that the animal was mobile. **(E)** Bin-averaged body directions (in deg, red arrow) and direction gradient (∂F/∂x) from 0 (blue) to 360 (yellow) deg for each given location across stimulus conditions (bin size = 7.54 cm/pixel; white bins indicate never visited locations).

**Supplemental Figure 6.**
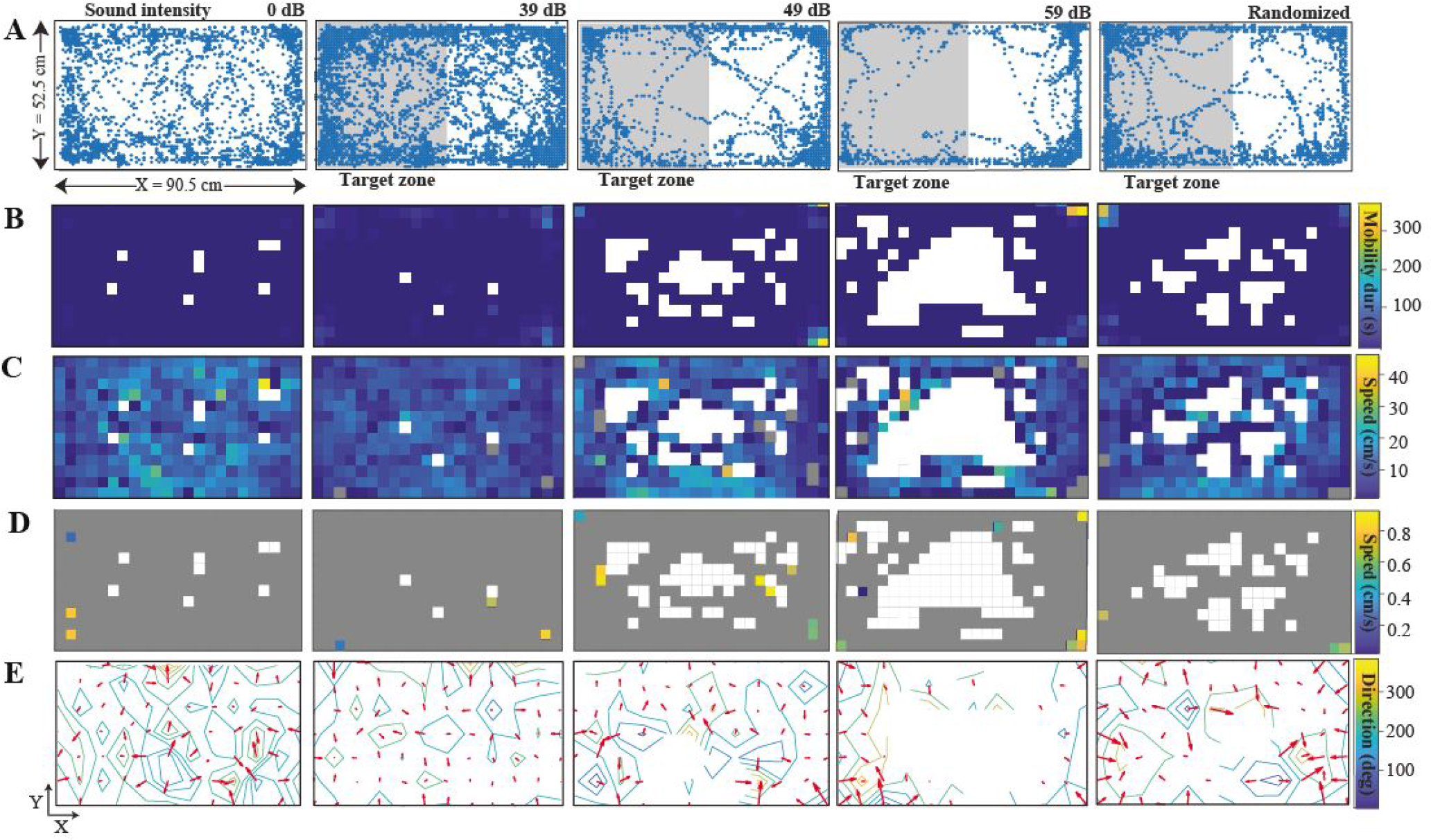
Locomotion activity in open field with discrete closed-loop feedback (animal 2). **(A)** Trajectory of animal 2 in open field arena for 5 different stimulus conditions: no feedback when the animal was in the target (T grey) zone (0 dB, session duration = 6 min, left panel), auditory feedback when the animal was in the T zone (39, 49, 59 dB discrete tone, middle 3 panels), or random feedback given pseudorandomly over time (3x 39, 49, 59 dB 10s tone, right panel). The position is the centre-of-mass of coincident multi-touches (bin size = 0.273 mm/pixel). **(B)** Bin-summed mobility duration (in s) for each given location in the open field (bin size = 80×80 pixels) across stimulus conditions. White bins indicate never visited positions. **(C)** The bin-averaged body speed (BS, in cm/s) for each given location when the animal was mobile (BS > 1 cm/s) across stimulus conditions. White bins indicate never visited places and grey bins indicate that the animal was immobile (BS < 1 cm/s). **(D)** Same as (C), but when the animal was immobile. Grey bins indicate that the animal was mobile. **(E)** Bin-averaged body directions (in deg, red arrow) and direction gradient (∂F/∂x) from 0 (blue) to 360 (yellow) deg for each given location across stimulus conditions (bin size = 7.54 cm/pixel; white bins indicate never visited locations).

**Supplemental Figure 7.**
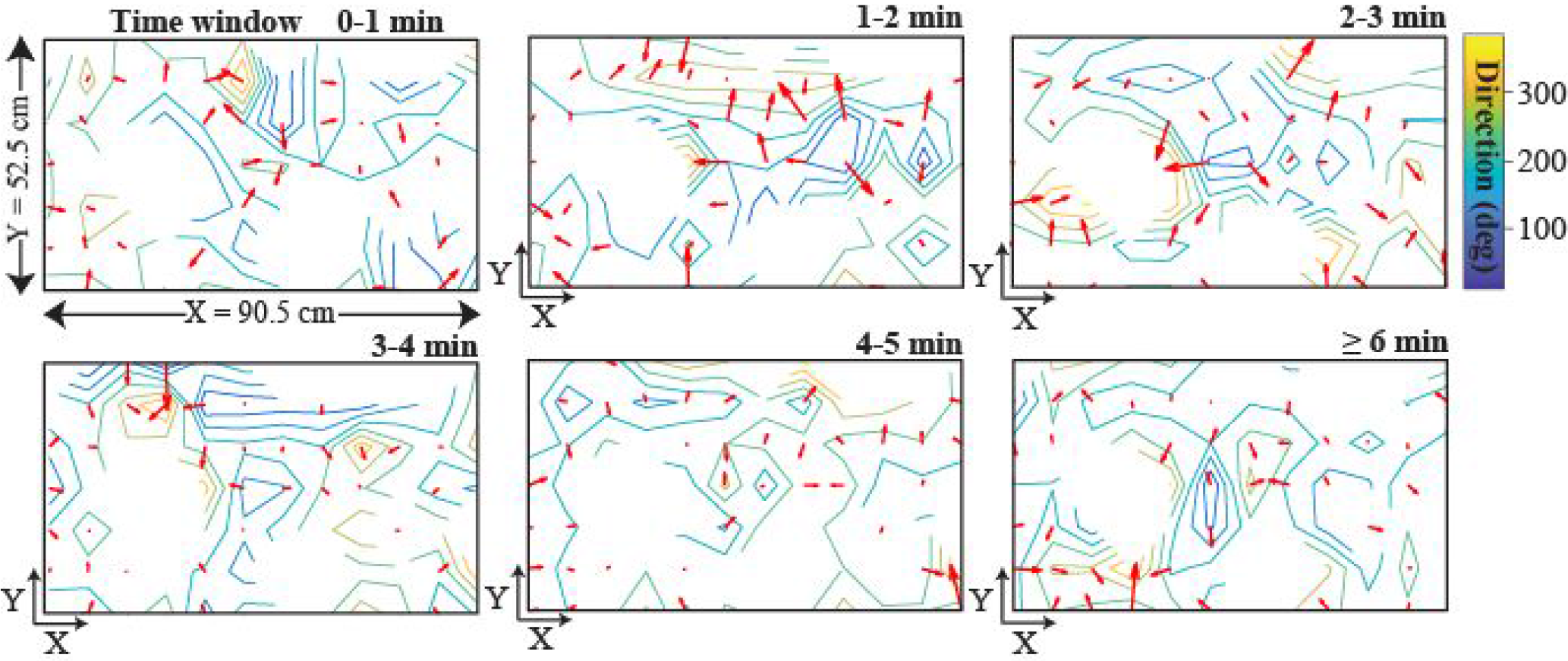
Vector map of body directions in open field for sequential time windows (animal 2). Vector map based on the bin-averaged body direction (red arrow) and direction gradient (∂F/∂x) from 0 to 360 deg of animal 2 during baseline conditions for subsequent 1-minute epochs (no feedback, session duration = 6 min, bin size = 7.54 cm/pixel, white bins indicate never visited locations).

**Supplemental Figure 8.**
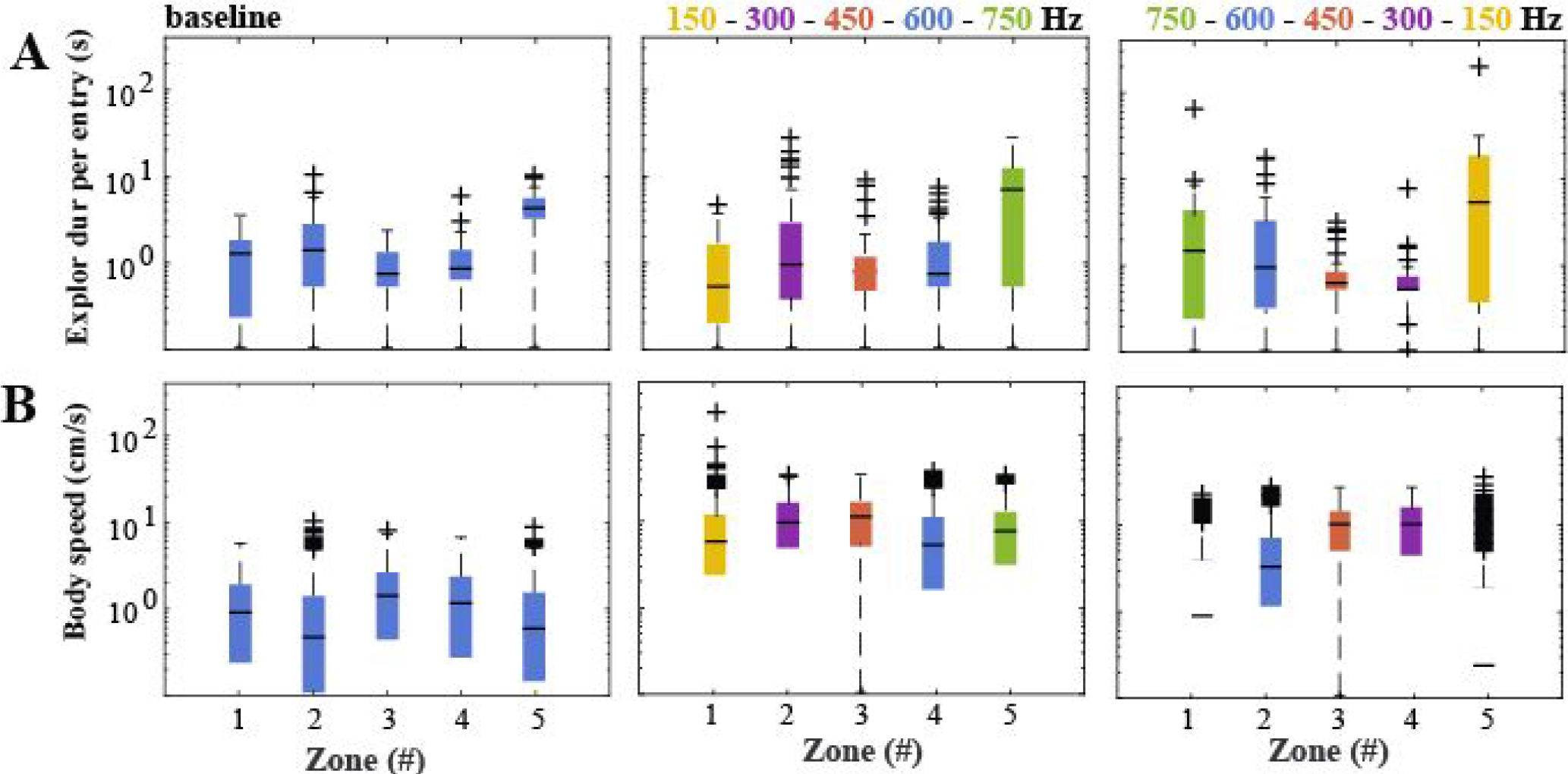
Locomotion activity in open field with continuous closed-loop feedback (animal 3). **(A)** Exploration duration per entry (ED, in s) of one animal to 5 virtual zones (z1-z5) in the open field for 3 stimulus conditions: no feedback (0 dB, baseline session, session period = 5 min, left panel), feedback given as a continuous sine tone with a frequency that increased (middle panel) or decreased (right panel) with the distance of the animal relative to zone 1 (ascending frequencies: 150, 300, 450, 600, 750 Hz, descending frequencies: 750, 600, 450, 300, 150 Hz, session period = 20 min/session). Boxplot with median ED (black line), 1st and 3rd quartile outliers (lower and upper lines), and outliers (cross symbols). Left panel: median ET_Z1_ = 1.2 s, 1st-3rd interquartile range (IQR_Z1_) = 0.2-1.7 s; median ET_Z2_ = 1.3 s, IQR_Z2_ = 0.5-2.6 s; median ET_Z3_ = 0.7 s, IQR_Z3_ = 0.5-1.2 s; median ET_Z4_ = 0.8 s, IQR_Z4_ = 0.6-1.3 s; median ET_Z5_ = 4.0 s, IQR_Z5_ = 3.1-5.1 s. Middle panel: median ET_Z1_ = 0.5 s, IQR_Z1_ = 0.2-1.6 s; median ET_Z2_ = 0.9 s, IQR_Z2_ = 0.4-2.8 s; median ET_Z3_ = 0.75 s, IQR_Z3_ = 0.5-1.2 s; median ET_Z4_ = 0.7 s, IQR_Z4_ = 0.5-1.6 s; median ET_Z5_ = 6.4 s, IQR_Z5_ = 0.5-11.4 s. Right panel: median ET_Z1_ = 1.6 s, IQR_Z1_ = 0-6.4 s; median ET_Z2_ = 5.4 s, IQR_Z2_ = 2.1-11.0 s; median ET_Z3_ = 15.3 s, IQR_Z3_ = 8.0-21.0 s; median ET_Z4_ = 15.3 s, IQR_Z4_ = 6.9-22.3 s; median ET_Z5_ = 0.47 s, IQR_Z5_ = 0-3.2 s. **(B)** Same as (A), but for the body speed (BS, in cm/s). Left panel: median BS_Z1_ = 10.2 cm/s, IQR_Z1_ = 4.4-16.2 cm/s; median ET_Z2_ = 6.8 cm/s, IQR_Z2_ = 2.7-13.3 cm/s; median ET_Z3_ = 13.5 cm/s, IQR_Z3_ = 6.5-19.8 cm/s; median ET_Z4_ = 11.8 cm/s, IQR_Z4_ = 4.8-18.5 cm/s; median ET_Z5_ = 7.8 cm/s, IQR_Z5_ = 3.2-14.1 cm/s. Middle panel: median ET_Z1_ = 7.1 cm/s, IQR_Z1_ = 2.9-12.7 cm/s; median ET_Z2_ = 4.8 cm/s, IQR_Z2_ = 1.3-10.6 cm/s; median ET_Z3_ = 10.7 cm/s, IQR_Z3_ = 5.0-17.1 cm/s; median ET_Z4_ = 9.1 cm/s, IQR_Z4_ = 4.5-15.5 cm/s; median ET_Z5_ = 5.4 cm/s, IQR_Z5_ = 2.0-10.7 cm/s. Right panel: median ET_Z1_ = 1.6 cm/s, IQR_Z1_ = 0-6.4 cm/s; median ET_Z2_ = 5.4 cm/s, IQR_Z2_ = 2.1-11.0 cm/s; median ET_Z3_ = 15.3 cm/s, IQR_Z3_ = 8.0-21.0 cm/s; median ET_Z4_ = 15.3 cm/s, IQR_Z4_ = 6.9-22.3 cm/s; median ET_Z5_ = 0.5s, IQR_Z5_ = 0-3.2 cm/s.

